# Single-nuclei histone modification profiling of the adult human central nervous system unveils epigenetic memory of developmental programs

**DOI:** 10.1101/2024.04.15.589512

**Authors:** Mukund Kabbe, Eneritz Agirre, Karl E. Carlström, Fabio Baldivia Pohl, Nicolas Ruffin, David van Bruggen, Mandy Meijer, Luise A. Seeker, Nadine Bestard-Cuche, Alex R. Lederer, Jilin Zhang, Virpi Ahola, Steven A. Goldman, Marek Bartosovic, Maja Jagodic, Anna Williams, Gonçalo Castelo-Branco

## Abstract

The adult human central nervous system (CNS) is remarkably complex, with neural cells displaying extensive transcriptional heterogeneity. However, how different layers of epigenetic regulation underpin this heterogeneity is poorly understood. Here, we profile the adult human CNS from distinct regions, for chromatin accessibility at the single-nuclei level. In addition, we simultaneously co-profiled the histone modifications H3K27me3 and H3K27ac at the single nuclei-level, providing their first map in all major human CNS cell types. We unveil primed chromatin signatures at HOX loci in spinal cord-derived human oligodendroglia (OLG) but not microglia. These signatures were reminiscent of developmental OLG but were decoupled from robust gene expression. Moreover, using high-resolution Micro-C, we show that induced pluripotent stem cell (iPS) derived human OLGs exhibit a HOX chromatin architecture compatible with the primed chromatin in adult OLGs, and bears a strong resemblance not only to OLG developmental architecture, but also high-grade pontine gliomas. Thus, adult OLG retain epigenetic memory from developmental states, which might enable them to promptly transcribe Hox genes, in contexts of regeneration, but also make them susceptible to gliomagenesis.

## Introduction

The human central nervous system (**CNS**) is a complex tissue encompassing the brain and spinal cord. It contains billions of diverse cells acting in concert to carry out executive functions including relay of sensory and motor input as well as integration, assimilation, and storage of information. The CNS arises from a uniform neural tube early during human development, which then undergoes waves of proliferation and transcription factor mediated patterning that ultimately leads to regionalization at the posterior-anterior and ventral-dorsal axis. This general patterning information is essential to determine the identity of the different areas in early stages in development, leading subsequently to the specification of unique neural types in the different regions, which themselves have distinct gene regulatory programmes in place.

Mature oligodendrocytes (**MOLs**) wrap neuronal axons with lipid-rich myelin, enabling rapid saltatory conduction of the action potential, consequently allowing precise coordination between different areas of the CNS^1–3^. MOLs are primarily found within the white matter (**WM**) areas of the CNS, whereas their progenitor population – oligodendrocyte precursor cells (**OPCs**) – are uniformly distributed throughout the CNS. In the mouse CNS, OPCs arise in distinct developmental waves, but transcriptionally converge after birth before differentiating to transcriptionally divergent MOLs in the adult mouse CNS.^3–5,6^ Previous studies investigating the oligodendroglial (**OLG**) lineage have identified region-specific transcriptomic differences in the human CNS ^7,8^.

While the transcriptome of neural cells in the human adult CNS has been well-characterized, the underlying regulatory chromatin landscape remains largely obscure. As the central repository of genetic information, chromatin is packaged inside the nucleus in a predominantly inaccessible state. Different regions of the genome are made accessible to allow for transcription and regulation of gene expression. Therefore, studying the accessible chromatin landscape provides a snapshot of the underlying regulatory blueprint defining a cell state^9–14^. Previous single-cell studies have leveraged this to identify human organ-specific regulatory elements at an organism level^15,16^, to profile the regulatory circuits underlying cell specification and differentiation during development^16,17^ and to understand how single-nucleotide variants disrupt regulatory element function in neurodegenerative diseases and cancers^18–23^. However, chromatin accessibility covers only an outer layer of epigenetic regulation. Post-translational modifications (**PTM**) at histones in nucleosomes and at the DNA have been shown to play essential roles in the regulation of transcription^24^. Delving a step further to capture nucleosomal information at the histone level has the potential to enhance the resolution by functionally annotating specific regions of the genome. The rise of powerful single-cell epigenomic technologies has now made this level of profiling attainable. Single-cell studies covering DNA methylation^25^ and individual histone modifications^26–36^ have started elucidating these epigenetic landscapes in the mouse adult CNS; however the exploration of the adult human CNS remains limited and currently only DNA methylation and chromatin architecture have been analysed at the single cell level^37^. Regarding histone modifications, current datasets are restricted to single cell analysis of H3K27me3 of a glioblastoma tumour from one patient^30^ and bulk characterization of glioblastoma tumours^18^, or specific fluorescent-activated nuclei sorted subpopulations in the human cortex^38^, while multi-modal single-cell profiling of the different cell states in the adult human CNS is still lacking.

Here, we provide a single-cell chromatin accessibility dataset of the adult human CNS across three regions spanning the anteriorposterior axis of the human CNS in different adult ages and sex, together with the first joint multi-modal single-cell histone PTM dataset in the adult human CNS. These resources, available at the UCSC Cell and Genome Browsers (https://cns-nanocuttag-atac.cells.ucsc.edu), allow us to define for the first time H3K27me3 and H3K27ac landscapes in the major CNS cell populations across regions, leading for instance to the identification of a novel enhancer for the human *SOX10* gene, which encodes a transcription factor essential for oligodendrogenesis. We also define neural cell-specific regulatory networks, identifying transcription factors that had not previously been associated with specific neural cell states. Importantly, we determine that the chromatin state of adult OLGs is reminiscent of their developmental counterparts, at the chromatin accessibility, histone PTM and chromatin architecture levels. Our analysis indicates that epigenetic memory of key developmental genes is in place in adult OLGs, which might prime these cells to rapidly activate transcription of Hox loci, and thus participate in regenerative processes, while making them susceptible to being hijacked in tumour transformation, such as gliomagenesis.

## Results

### Single-nucleus ATAC-seq of the adult human CNS reveals differential chromatin and TF motif accessibility in different CNS cell types

We collected a cohort of 60 tissue samples from 20 post-mortem donors ranging in age from 34 to 74 years old and with equal representation of both sexes. From each donor, we had frozen tissue samples from three distinct regions of the CNS: primary motor cortex (**BA4**), cerebellum (**CB**) and cervical spinal cord (**CSC**) (**Supplementary Table 1**). Based on tissue quality metrics^39^ (**see Methods**), we isolated WM-dominant areas from the tissue, dissociated them into single-nuclei suspensions and performed single nucleus ATAC-seq (**snATAC-seq**) using the 10x Genomics Chromium platform (**Fig. 1a**). After sequencing and stringent quality control based on the number of unique reads and per-cell TSS enrichment score (**Extended Data Fig. 1**), we retained 108,626 nuclei representing all three regions (Motor Cortex: 55037 cells, Cerebellum: 34819 cells, Cervical Spinal Cord: 18770 cells), with a median of 8154 fragments per cell, and a median TSS enrichment score of 10.7 (**Extended Data Fig. 1a-b**). We built a count matrix using a 2kb binned genome as the features and retained the top 100,000 most accessible bins^40,41^. After Term Frequency-Inverse Document Frequency (**TF-IDF**) normalization, dimensionality reduction and clustering on the nearest neighbour graph, we obtained 16 distinct clusters across the three regions (**Fig. 1b,d**). Since gene expression is correlated to promoter and TSS activity, we built a gene activity matrix using the aggregate signal spanning gene promoters and used the signal as a proxy for gene expression. To annotate the cell types, we assigned a metagene score for the broad cell types found in the CNS^29^ (**see Methods**). Using metagene scores, we identified all major cell types including cerebellar excitatory neurons (**CBEX**: 21801 cells), cerebellar inhibitory neurons (**CBINH**: 1301 cells), cortical excitatory neurons (**CXEX**: 9671 cells), cortical inhibitory neurons (**CXINH**: 4353 cells). Conversely, the glial cell populations were relatively homogeneous, and we identified mature oligodendrocytes (MOL: 45325 cells), OPCs (OPC: 5130 cells), microglia (**MIGL**: 9528 cells), astrocytes (**AST**: 10098 cells) and pericytes/endothelial cells (**ENDO**: 1419 cells) (**Extended Data Fig. 2b-c**). We identified several marker genes for the different populations including *SOX10* for the OLG lineage, *PDGFRA* for OPCs, *PLP1* for MOLs, *AQP4* for AST, *AIF1* for MIGL, *RBFOX3* for excitatory neurons and *GAD1* for inhibitory neurons (**Fig. 1d-e; Extended Data Fig. 2c**). We confirmed the validity of our metagene-derived annotations by integrating the gene activity object with a paired single-nuclei transcriptomic dataset of the same cohort^39^ and a previously published dataset^42^ (**Extended Data Fig. 2a**).

**Fig. 1.**
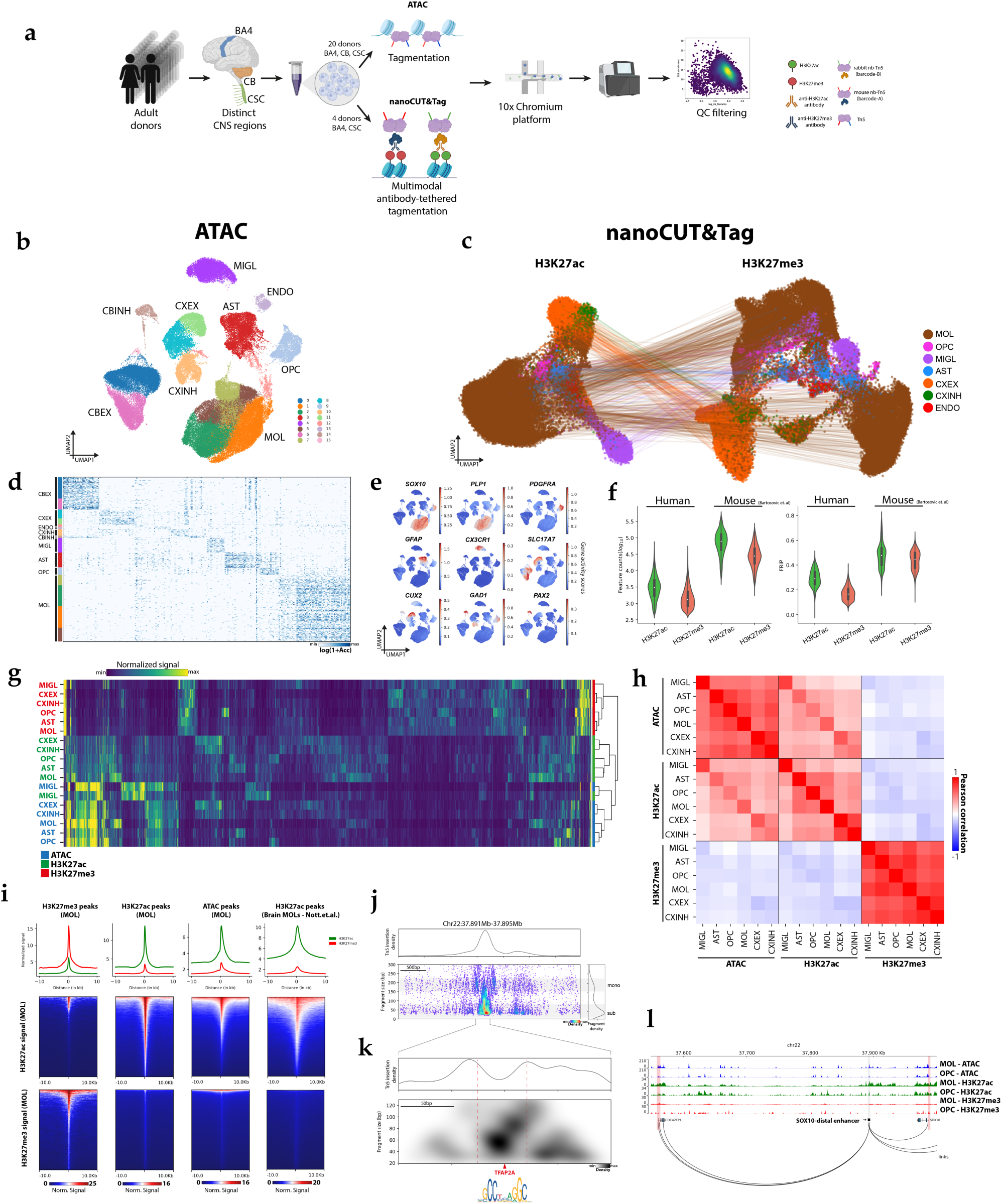
a. Schematic for snATAC-seq and nanoCUT&Tag experiments in adult human tissue. b. 2D Uniform manifold approximation and projection (UMAP) of the ATAC dataset coloured by clusters and labelled by cell type. c. 2D Uniform manifold approximation and projection (UMAP) of the nanoCUT&Tag dataset from both modifications. Coloured lines connect the same cells in both modalities. d. Heatmap showing differentially accessible peaks across different clusters and cell types. e. Gene activity scores for different genes in the identified cell types. f. Quality metric violin plots showing the number of unique features (left) and fraction of reads in peaks (FRiP) for H3K27ac (green) and H3K27me3 (red) in this dataset compared to those in a previously published dataset in mouse (Bartosovic et.al. 2022). g. Trimodal clustering of the genome highlights patterns of signal distribution across all cell types. h. Correlation matrix of ATAC, H3K27ac and H3K27me3 signal in each cell type shows strong correlation between active marks for individual cell types, and anti-correlation with the repressive H3K27me3. i. Meta-signal enrichment plots for H3K27ac (green) and H3K27me3 (red) in the MOL population. Top row: Line plots showing signal enrichment for the two modalities at different peak sets. Middle and bottom row: Heatmap showing H3K27ac (middle) and H3K27me3 (bottom) signal enrichment across different peak sets. Peak sets (left to right): H3K27me3 peaks, H3K27ac peaks, ATAC peaks, ATAC peaks from a previously published dataset (Nott et.al. 2019). j. V plot showing tagmentation pattern and density in a 3.5kb locus around the SOX10-distal enhancer. Top: Density plot showing density of the Tn5 insertion events, seen enriched at the site of the enhancer. Bottom right: Fragment size distribution, with sub-nucleosomal and mono-nucleosomal bands visible. Bottom left: Scatter plot of fragments. Dots represent mid-points of fragments in the OLG lineage and coloured by density of fragments. k. 330bp locus zoom in at the enhancer site. TF footprint is seen as dip in the Tn5 insertion frequency (top) and is marked by a red dotted line. V plot shows density of fragments at the centre of the footprint. A TFAP2A motif was found in the centre of the footprint. l. Genome browser tracks the *CDC42EP1*-Enhancer-*SOX10* locus showing ATAC, H3K27ac and H3K27me3 signal in MOLs and OPCs and cicero links.

We then investigated transcription factor (**TF**) motif accessibility differences in the different cell types in our dataset, using chromVar^13^. Clustering on the motif deviations identified marker TFs for the different populations, for instance SOX transcription factors for oligodendroglia, or a specific RORA enrichment in the cerebellar neurons, which is required for cerebellar Purkinje cell maturation^43^ (**Extended Data Fig. 2d-e**). Interestingly, while the neurons clustered distinctly according to region and broad electrophysiological profiles, we did not observe the same distinctions within the glial populations (**Extended Data Fig. 2f**), suggesting the chromatin states for glial cells may be more region-agnostic and present plasticity, to account for their varied functions.

### Single nucleus nanoCUT&Tag H3K27ac and H3K27me3 profiling of major cell populations in the adult human CNS

Specific histone PTMs are associated with active and repressed transcriptional states^24,44–46^. Histone PTMs serve as signalling beacons on the chromatin and can inform about the functional state of local chromatin, providing a more granular resolution than accessibility alone. We have recently developed nanoCUT&Tag, which allows targeting of two histone PTMs simultaneously in the same cell using uniquely barcoded nanobody-Tn5 (**nanoTn5**) fusion proteins^31^. While nanoCUT&Tag has been applied at a single cell level to the mouse brain^31^, it has not been applied to the human CNS. For this purpose, we adapted the protocol of nuclear extraction performed for the archival tissue in snATAC-seq for nanoCUT&Tag (**see Methods**). We were able to successfully profile H3K27ac (active mark) and H3K27me3 (repressive mark) simultaneously, in three cervical spinal cord and three cortical frozen archival tissue samples from a total of four donors (**Fig. 1a, c**). The custom Tn5 barcodes allows us to de-multiplex the data into the respective modalities and proceed with individual processing, while the shared 10x barcode incorporated in the GEMs links the cells from each modality together. After using our custom demultiplexing and cell-calling pipeline^31^ (**Extended Data Fig. 3a-c; see Methods**), we identified 66,113 and 66,727 barcodes in the

H3K27ac and H3K27me3 datasets respectively, with an 88% barcode overlap, leading to 58,696 shared cells. We captured a median of 2099 and 1268 unique fragments for H3K27ac and H3K27me3 respectively, lower but still comparable to metrics in our published mouse datasets^31^ (**Fig. 1f**). After calling peaks, the fraction of reads in peaks (FRiP) was 0.29 and 0.17 for H3K27ac and H3K27me3, respectively (**Fig. 1f**). The low FRiP values are expected since our method relies on two-step tagmentation leading to variable fragment length (**see Methods**). Nonetheless, we have shown previously that lower FRiP does not negatively impact clustering and cell type identification^31^.

H3K27ac is an active mark and canonically marks active enhancers and promoters^44^. As such, the signal has strong correlation with chromatin accessibility. Hence, we integrated the H3K27ac dataset with our snATAC-seq dataset (**Extended Data Fig. 3d; see Methods**). After integration and label transfer, we identified the MOL (40268 cells), CXEX (8821 cells), MIGL (4297 cells), AST (2169 cells), CXINH (1774 cells), OPC (820 cells) and ENDO (547 cells) populations (**Extended Data Fig. 3d-e**). Genome browser tracks showed the expected enrichment of the respective signal for each of the identified cell types (**Extended Data Fig. 3f**). To ascertain the quality of cell-type specific histone PTM landscapes, we then clustered all genes based on the joint H3K27ac and H3K27me3 profile in each cell type. This allowed the identification of distinct clusters of genes, with increased active or repressive mark. We then performed a cell-type enrichment analysis using the genes in the active-mark dominated clusters and confirmed that the same cell types that were queried were enriched in the respective populations (**Extended Data Fig. 4**).

To check the specificity of the antibody signal, we generated meta-signal plots for both modalities looking at H3K27ac and H3K27me3 peaks in MOLs. We observed a strong enrichment of each mark’s signal in the respective peak set with minimal cross-mapping, confirming the specificity of the H3K27me3 and H3K27ac signal. The small cohort of genes showing overlap in H3K27ac and H3K27me3 signal might correspond to poised genes^47^. We also confirmed the enrichment of H3K27ac signal and absence of H3K27me3 at peaks called in the ATAC data, and in a published bulk H3K27ac dataset^38^ (**Fig. 1i**). We were unable to find a representative H3K27me3 dataset in the human brain to benchmark against, highlighting this as a unique dataset providing both H3K27ac and H3K27me3 at single-cell resolution in the brain and spinal cord.

We then used the ATAC, H3K27ac and H3K27me3 human CNS single-nuclei datasets to perform trimodal clustering of the entire genome and identify patterns and correlations between the three modalities in neural cell types (**Fig. 1g; see Methods**). Although we identified expected patterns of the three signals across the genome, the overall matrix looked scattered, highlighting the complexity of the regulatory genome, but also the sparsity of the datasets. We identified different regions of the genome that were 1) accessible in all cell types, likely corresponding to housekeeping genes; 2) specifically accessible in each cell type with corresponding H3K27ac signal; 3) presented shared H3K27me3 in all cell types (**Fig. 1g**). We then used the signal in all genomic bins to look at correlations between the different cell types, and observed, as expected strong anti-correlation between the active ATAC and H3K27ac marks with the inactive H3K27me3 mark (**Fig. 1h**). Interestingly, chromatin accessibility exhibited a higher correlation across the different cell types overall, unlike H3K27ac, which presented higher correlation between glial populations or between neuronal populations (**Fig. 1h**). These findings suggest that H3K27ac may be a better discriminant of cell type-specific regulatory activity compared to chromatin accessibility.

### Identification of a novel enhancer for the *SOX10* gene in human OLGs

Promoters and enhancers are characterized by open chromatin and H3K27ac deposition, which makes our single-nuclei ATAC and nanoCUT&Tag datasets well suited to identify known but also novel candidate cis-regulatory elements (**cCREs**) (**Extended Data Fig. 5a; Supplementary Table 2; see Methods**). To determine if we indeed could identify known CREs, we focused on the *SOX10* gene, which has several well characterized enhancers, some of which are operational in the OLG lineage^48^. Indeed, we found that the *SOX10* promoter was co-accessible with several peaks corresponding to the enhancers in OLG, but not in other lineages^49^ (**Extended Data Fig. 5b,d**). Importantly, we found that the promoter was also co-accessible with two distal upstream peaks, previously not associated with *SOX10* (**Extended Data Fig. 5b)**. One peak corresponded to the promoter of the *ANKRD54* gene, which we found to be accessible in all cell types. However, the second peak, situated 91kb upstream of *SOX10* was intergenic, and accessible only within the OLG lineage, suggesting it may be an uncharacterized cell-type specific enhancer for this lineage.

Interestingly, we also found that this *SOX10* distal enhancer was co-accessible with the *CDC42EP1* promoter, located 300kb away (**Extended Data Fig. 5b**). *CDC42EP1* encodes the effector protein of CDC42, which is associated with myelin sheath compaction in MOLs^50^. Interestingly, we observed increased accessibility of the *CDC42EP1* promoter specifically within MOLs, but not OPCs. We checked the co-accessibility links within the OPCs and MOLs separately, and found *SOX10* interactions with the canonical enhancers, and the new enhancers in both populations, but the CDC42EP1-enhancer connection only in MOLs (**Extended Data Fig. 5e**), suggesting that the chromatin looping concerning this enhancer is altered upon OLG lineage progression. We also queried this locus in our nanoCUT&Tag dataset and observed an increase in the H3K27ac signal at the identified enhancer in MOLs and OPCs, with an increase in H3K27ac at the *CDC42EP1* promoter in MOLs specifically (**Fig. 1l**).

This enhancer has high conservation with four other species including the phylogenetically close species rhesus monkey and mouse (**Extended Data Figure 5c**). The PhyloP score for the bases in this locus, which measures evolutionary conservation^51^, was also positive and higher than the negatively scored flanking bases. This indicates slower mutation rates, and therefore higher likelihood of conservation, which is characteristic of enhancer evolution^52^. We used V plots^53,54^ to visualize the density and pattern of tagmentation events in a 3.5 kb locus spanning the enhancer. The plot revealed a strong density of fragments corresponding to the sub and mono-nucleosomal bands, but also intermediate-length fragments, indicative of dynamic TF-bound open chromatin (**Fig. 1j**). Zooming in further into a 300bp window revealed a clear TF footprint in the tagmentation density, corroborated by the expected fragment distribution in the V plot (**Fig. 1k**). A motif analysis of the core footprint identified a TFAP2A motif coinciding well with the expected binding site from the V plot. The AP2-alpha TF has been shown to regulate *SOX10* expression but specifically via the U3 enhancer^48^, which is distinct from this distal enhancer. This suggests that a new distal enhancer may be regulating *SOX10* expression in the OLG lineage.

Collectively, these findings propose a new enhancer that regulates *SOX10* in both MOLs and OPCs, and selectively regulates *CDC42EP1* in MOLs and highlight the potential of the ATAC and nanoCUT&Tag datasets in identifying and functionally annotating putative regulatory elements.

### Novel core regulatory TF networks in adult human neural cell types

Using our H3K27ac and ATAC datasets, we constructed a core TF regulatory network for each identified cell type, by looking at enhancer and TF motif accessibility. In the constructed TF networks, we looked at nodes with more outgoing connections than incoming connections, indicating these were strong regulators in the TF network (**Fig. 2a**). Along with the expected TFs such as OLIG2 in MOLs and OPCs, IRF2 in MIGL, CUX2 and SATB1 in cortical neurons (CXEX and CXINH), we also identified new regulators, such as OLIG3 in MIGL and ZBTB38 and HMX1 in AST (**Fig. 2b; Supplementary Table 3**), highlighting the utility of this dataset in identifying novel TF networks in neural cell types. PAX3 was identified in OPCs, which could reflect a dorsal origin of a subset of OPCs^18^. We also identified within the OPC population, but not in MOLs, PRRX1, a TF shown to regulate OPC quiescence^55^. Interestingly, when we looked at expression levels of the core TFs, we found that the highest expressed TF in the network was not the strongest TF, except for CAMTA1 in CXINH neurons. Indeed, ZBTB20 which is upregulated after cerebral ischemia in neural progenitor cells^56^, was the highest expressed TF in the AST, MOL and OPC populations but was only the 9^th^, 20^th^ and 29^th^ strongest TF in the respective networks (**Fig. 2c-d**).

**Fig.2.**
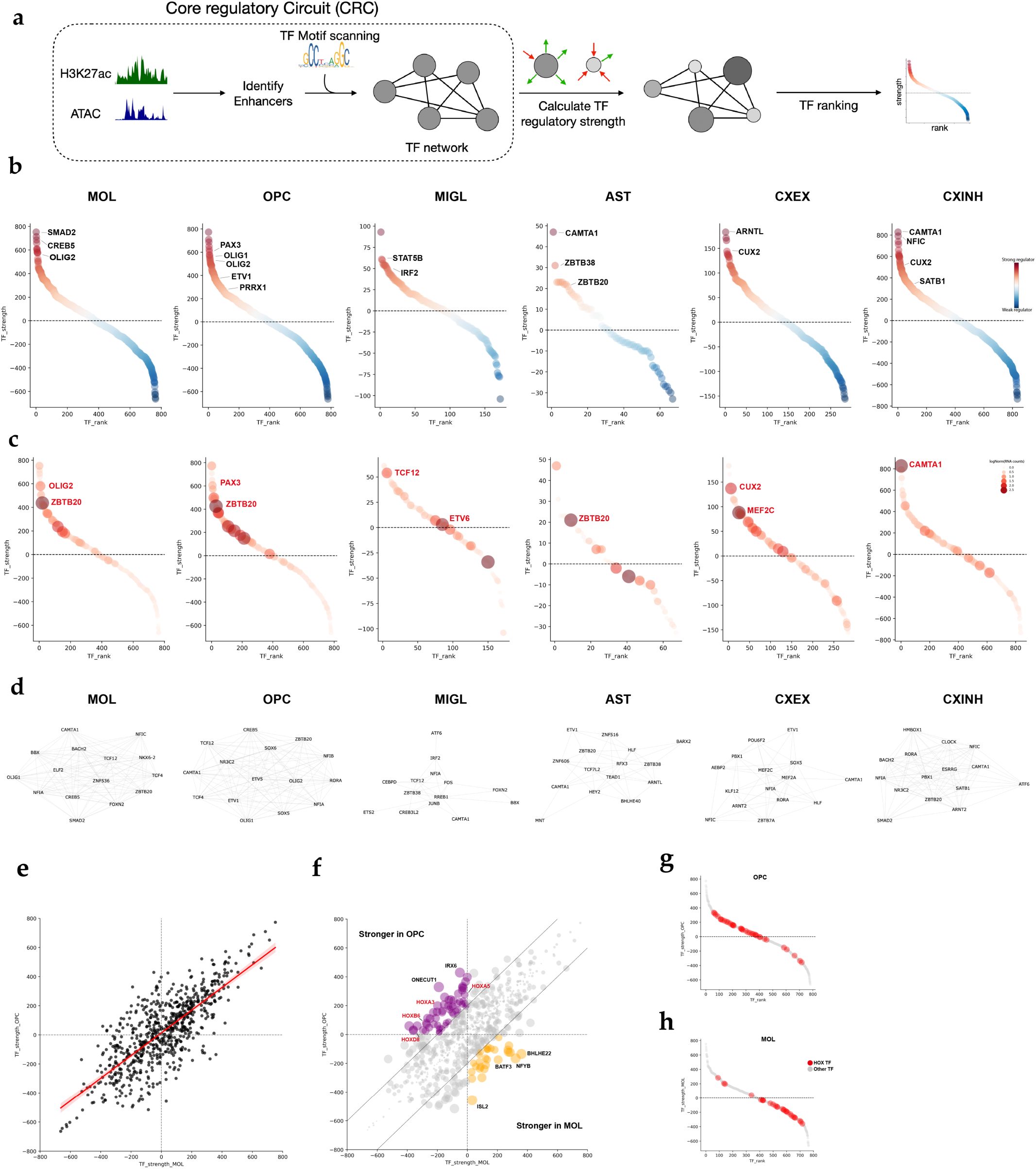
a. Schematic for constructing the regulatory TF circuit and assigning a regulatory strength based on the difference in the incoming and outgoing connections for each TF node in the network. b. Ranked scatter plot showing the strength of core transcription factors (TF) identified within the regulatory network for different cell types. Strength is measured by the difference in number of outgoing connections (out degree) and incoming connections (in degree). Top TFs are marked for each population. c. Same plot as in a) but size and colour intensity of each dot represents the average (log normalized) expression of that TF in the population. The TF with the highest expression in each network is shown in red. d. Network of the top 15 TFs (by expression) in each cell type. Density of edges in the network reflect the correlation between the TF strength and the expression of the TF. e. Scatterplot showing the strength of the shared TFs in the OPC and MOL core networks. Regression line is shown with a 95% CI. f. Same as in d) but size of the dots represents the difference in the strength of the TF in the OPC and MOL populations. TFs stronger in OPC and MOL are shown in purple and orange respectively. A subset of the HOX TFs is marked in red. g. Ranked scatterplot highlighting the position of the HOX TFs in the OPC population. h. Same as f) but in MOL population.

We also investigated whether TF networks change upon cell type differentiation using OLG as a paradigm, and specifically the OPC and MOL populations. Overall, we found over 90% overlap in the TFs in each population’s core network, suggesting high TF coherence throughout the lineage (**Fig. 2e**). Nevertheless, we found several non-overlapping TFs with high regulatory strength and expression that might regulate the transition from OPCs to terminally differentiated MOLs, such as – ARX, MYC, SIX1 and FOXF2 in OPCs, and NFAT5, ARNT, ZNF566 and ZNF333 in MOLs (**Supplementary Table 3**).

We then looked at TFs that switched status from being a strong regulator in the progenitor state to a weaker regulator in the differentiated state (positive score in OPCs and negative in MOLs), or vice-versa, and found among others, BHLHE22 as a strong TF in the MOLs (**Fig. 2f**). *BHLHE22* expression has been shown to increase in OPCs upon T3 stimulation and plays a role in differentiation and myelination^57^, in line with our finding that BHLHE22 increases in strength in MOLs. In OPCs, we found ONECUT1, which regulates *NKX6-2* expression, a key TF in OPCs. Interestingly, we also found several TFs in the HOX family of proteins, as strong regulators in OPCs, with weaker strength in MOLs (**Fig. 2f; Extended Data Fig. 6a**). Though the regulator strength was higher in OPCs, they were also present in the MOL core network, suggesting they may have higher potential in the OPC stage, which then goes down upon differentiation (**Fig. 2g-h**). While the activity reflects the presence of spinal cord OPCs, these HOX TFs were not identified in the core network for any of the other cell types, suggesting this may be specific to the adult OLG lineage (**Supplementary Table 3**).

### Spinal cord adult OLGs exhibit increased accessibility at HOX genes, decoupled from gene expression

The HOX family of proteins are evolutionarily conserved transcription factors expressed in embryonic development for patterning^58^. Since they are expressed within the developing spinal cord^59^, and we observed potential regulatory activity in human adult OLG, we investigated if there was regional specificity to these TFs, by looking at differential chromatin accessibility (**DA**) between the motor cortex (BA4) and cervical spinal cord (CSC)-derived OLGs. We found differential accessibility in MOLs and OPCs at the promoter/gene body of genes that were previously observed to be differentially expressed in these regions, such as *PAX3*, *SKAP2*, *SPARC*, *HCN2* in CSC-OLGs and *FOXG1*, *NELL1* in BA4-OLGs^39^ (**Fig. 3a**). Moreover, we also observed several HOX cluster genes presenting differential chromatin accessibility in the cervical spinal cord-OLGs when compared to cortical OLGs (**Fig. 3a-b, Extended Data Fig. 6b**). Genome browser tracks further revealed that both the OLG lineage and AST, but not MIGL presented increased accessibility at the HOX genes in spinal cord (**Fig. 3c**). The pattern of HOX gene accessibility that we observed was in line with the genes that would be expressed during development in the cervical area of the spinal cord^60^.As promoter activity can serve as a proxy for transcriptional activity, we reasoned that the promoter chromatin accessibility signal may correspond to these genes being expressed. We thus co-profiled both chromatin accessibility and gene expression in the same cell using the 10x Genomics Chromium multiOme platform on samples from the motor cortex and the cervical spinal cord (**Extended Data Fig. 6c**). We integrated the multiOme-ATAC with our snATAC-seq dataset and annotated the cells using a k-nearest neighbours (kNN) classifier (**see Methods**). The co-profiled dataset revealed that chromatin accessibility and expression of genes associated with OLG identity, such as *PTPRZ1* (OPCs), *PLP1* (MOLs) and the TF-encoding *SOX10*, *OLIG1*, *OLIG2* genes were highly correlated (**Fig. 3d**). In contrast, we observed increased accessibility of several HOX genes in the cervical spinal cord-OLGs, while RNA expression was residual (**Fig. 3d**). To ascertain if the weaker RNA signal detected was a limitation of sequencing depth, we examined the expression of HOX genes in a high-depth single-cell transcriptomic atlas^61^ of the adult human brain. We observed similar residual levels of HOX gene expression (**Extended Data Fig. 6d**), suggesting a transcriptional and epigenomic decoupling at HOX loci in adult OLGs, unlike during development.

**Fig.3.**
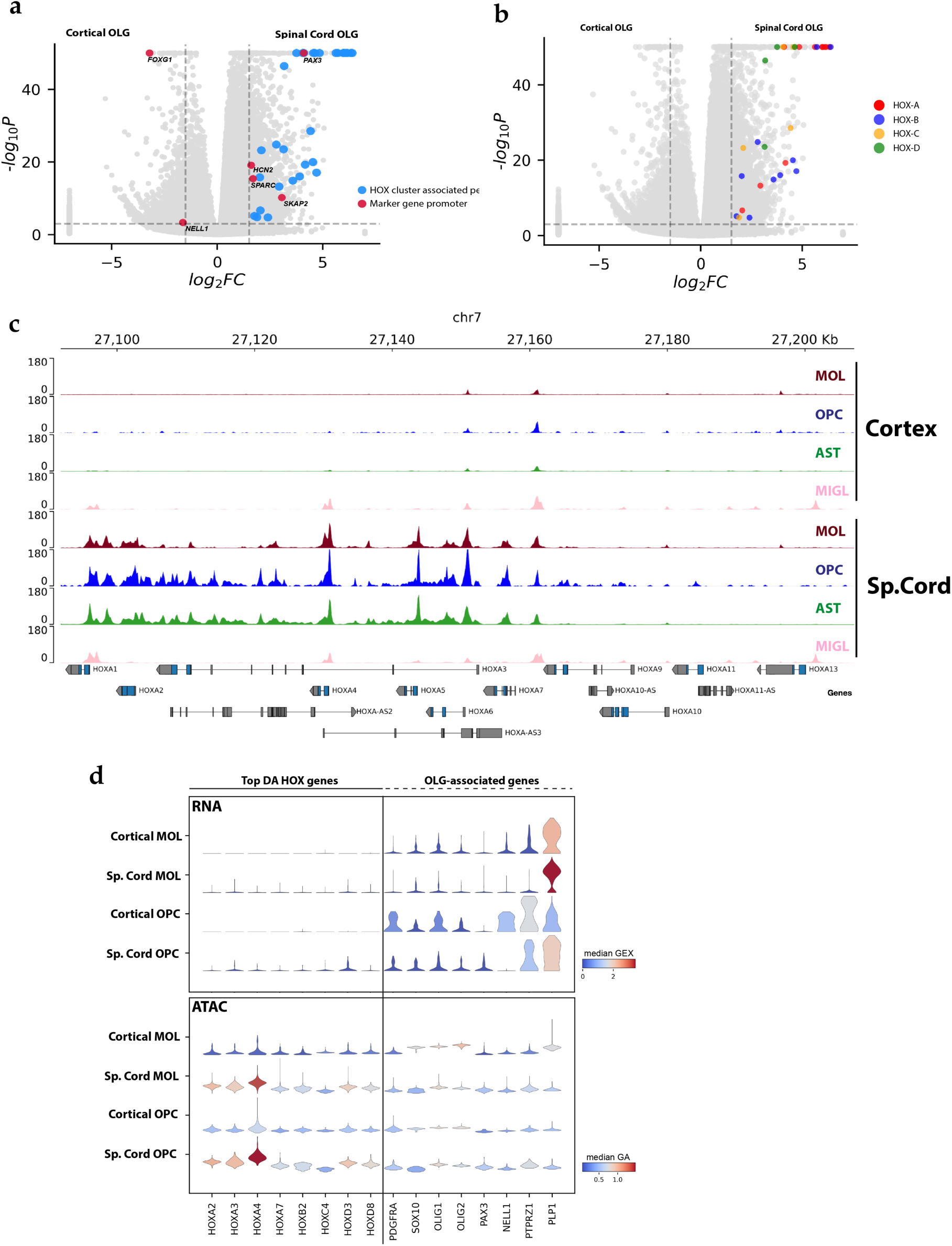
a. Volcano plot showing differentially accessible peaks in Spinal cord OLGs and cortical OLGs. Previously characterized marker genes are shown in red and labelled. HOX cluster-associated peaks shown in blue. Thresholds: adjusted P-value: 0.001, logFC = 1.5. b. Same as in a) but highlighting the specific HOX clusters identified as being differentially accessible. c. Genome browser snapshot showing the chromatin accessibility signal in AST, MIGL, MOL and OPC populations from the cervical spinal cord and motor cortex at the HOXA locus. MIGL signal is depleted in both regions, whereas AST, OPC and MOL exhibit accessibility in spinal cord. d. Stacked violin plots showing the expression (upper panel) and promoter accessibility (lower panel) in cortex and spinal cord derived MOLs and OPCs from a multiOme experiment. Most differentially accessible HOX genes and OLG marker genes are shown.

### HOXA/D genes are primed for expression in subsets of spinal cord-derived adult human oligodendroglia

Since HOX genes present open chromatin in adult human OLGs, while their expression is attenuated, mechanisms other than chromatin accessibility may control HOX gene transcription. We therefore checked for deposition of H3K27ac and H3K27me3 between spinal cord-OLGs and cortical-OLGs at these loci. Pseudo-bulk genome browser tracks showed that cortical cells displayed a pan cluster deposition of H3K27me3 reflecting the canonical pattern of Polycomb Repressive Complex 2 (**PRC2**) mediated HOX gene repression^62^ (**Fig. 4a-b**; **Extended Data Fig. 7a-b**). In contrast, our data indicated anti-correlated gradients of H3K27me3 and H3K27ac across the HOXA and HOXD clusters in spinal cord-OLGs (**Fig. 4a-b**). Moreover, we observed differential deposition of these marks along HOX genes, reminiscent of the concept of HOX gene collinearity during development^60,62,63^. The 3’ end of the clusters (for example, HOXA1 to HOXA7) showed elevated active marks (ATAC, H3K27ac) and a reduction of H3K27me3, while the 5’ end of the clusters (for example, HOXA10 to HOXA13) showed the opposite with an almost sharp inversion in the middle of the cluster. Thus, the pattern observed in the human adult spinal cord-OLGs reflects a pattern that would be seen during embryonic development in the cervical regions of the spinal cord, when these HOX genes are actively transcribed, suggesting epigenetic memory of the developmental chromatin state in these adult OLGs.

**Fig.4.**
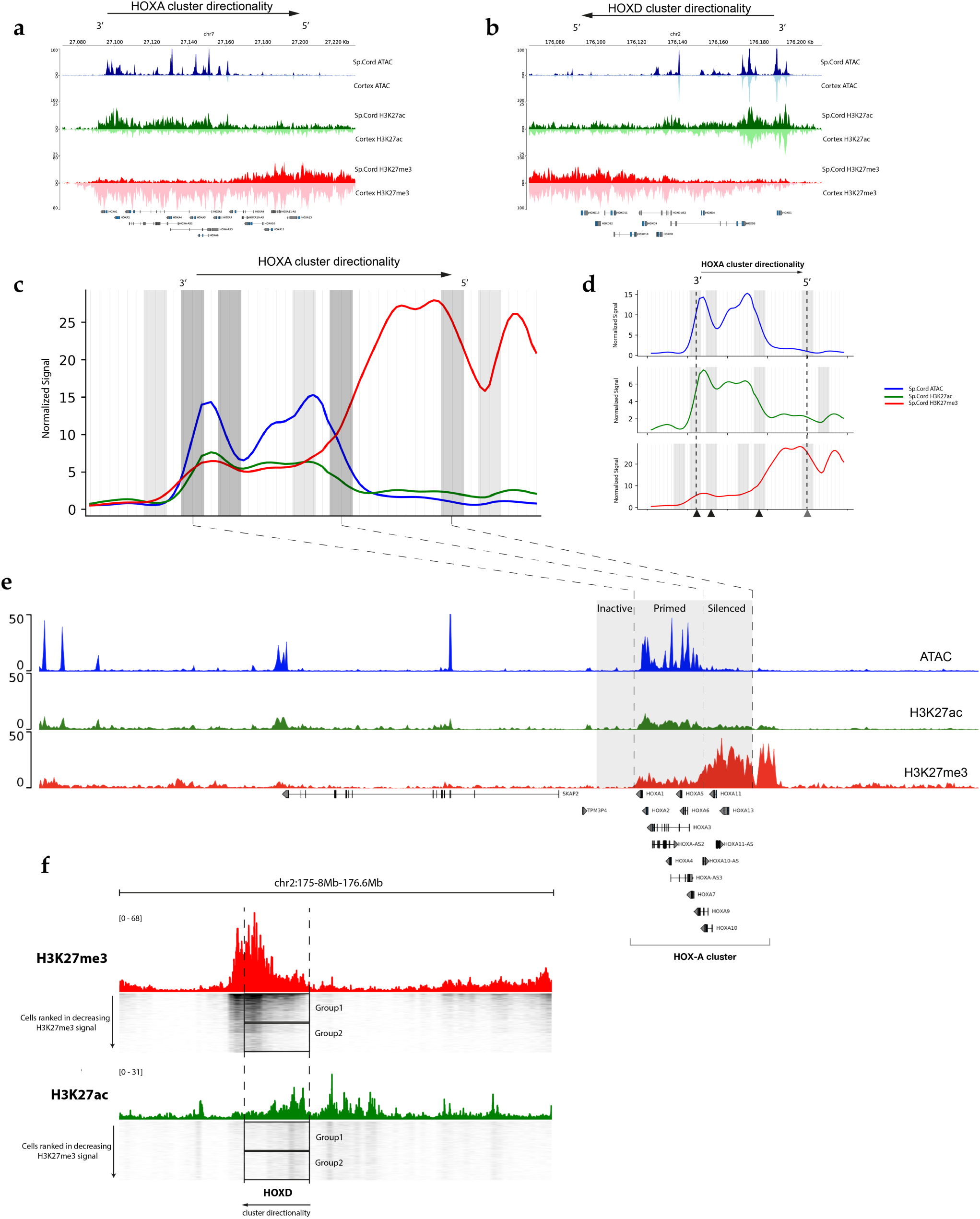
a. Genome browser tracks showing H3K27ac, H3K27me3 and ATAC signal in OLGs at the HOX-A in cervical spinal cord (upright track, darker shade) and motor cortex (inverted track, lighter shade). b. Same as in a) but for the HOX-D locus. c. Gaussian smoothed normalized signal from ATAC (blue), H3K27ac (green) and H3K27me3 (red) across the HOXA cluster with a 50kb flanking region upstream and downstream. Gray bars show the locations of the cumulative “signal boundaries” identified in each modality. Colour intensity reflects the cumulative signal boundary strength. d. Same as in panel c) but with each modality separated out. HOXA directionality is shown at the top, and arrows beneath show the medium (2 modalities) and strong (3 modalities) signal boundaries. e. Genome browser track of the HOXA cluster showing the location of the strong signal boundaries and the corresponding inactive, primed and silenced chromatin domains. f. Genome browser track around the HOXD locus (marked with dotted lines) with H3K27me3 (red) and H3K27ac (green) pseudobulk signal in spinal cord OLGs. Single cell tracks are shown below and sorted in order of decreasing H3K27me3 signal. Group 1 cells exhibit moderate H3K27me3 at the 3’ end while Group 2 cells show a depletion of the H3K27me3, while the amount of H3K27ac remains the same in both groups, suggesting Group2 cells may be expressing low levels of HOX genes.

To identify domains with significantly differential signal, we looked at the signal from all three modalities, in a genomic window covering the HOX clusters and spanning 50kb upstream and downstream of each cluster. We binned the region into discrete windows and identified borders between adjacent windows with significant changes in signal (**see Methods**). By overlapping the borders detected in each modality, we identified distinct domains across which there were changes in the levels of all three modalities. Within the HOXA cluster we identified three strong, one moderate border and one weak border (**Fig. 4c-d**). Two of the strong borders were situated near the 3’ end of the cluster, flanking HOXA1 to HOXA4, and were present in all modalities (**Fig. 4d-e**). A weak border around HOXA7 demarcated increased levels of H3K27me3 with increased chromatin accessibility, while a strong border at HOXA10 coincided with the point at which we saw the signal switch from predominantly accessible to predominantly repressed via H3K27me3. A moderate border marked the decrease of the heavily inactive signature at the 5’ end of the cluster (after HOXA13). The strong border identified at the 3’ end of the cluster suggested further nuance. Although the levels of H3K27me3 at the 3’ end (HOXA1 to HOXA7) were far lower than the 5’ end, it was distinctly greater than the flanking chromatin immediately upstream of the cluster and to the left of the identified border. Thus, we could demarcate three regulatory domains around the HOXA cluster – 1) inactive chromatin upstream of the cluster; 2) primed chromatin at the 3’ end and 3) silenced chromatin at the 5’ end (**Fig. 4e**). While the HOXB and HOXC clusters displayed moderate borders at the 3’ end of the respective clusters, the HOXD cluster displayed strong borders at the 3’ end and in the middle of the cluster, as in the HOXA cluster (**Extended Data Fig. 7c**). Thus, this indicates three distinct levels of the H3K27me3 repressive mark at HOX loci in adult OLGs, suggesting that this increased level of H3K727me3 at the 3’ end might be sufficient to prevent gene expression and maintain the genes in a primed state.

The multimodal nature of the nanoCUT&Tag data allows to look at the deposition pattern of H3K27ac and H3K27me3 within the same cell. We looked at the HOXD cluster and ranked the cells in decreasing order of H3K27me3 signal. At the 5’ end of the HOXD cluster (HOXD8 to HOXD13), we saw a clear pattern of high H3K27me3 signal in all cells. However, at the 3’ end we identified 2 sub-groups, displaying either a medium (Group1) or low (Group2) level of H3K27me3. Interestingly, when looking at the corresponding H3K27ac signal in the same cells, we did not see this bimodal distribution of H3K27ac at the 3’ end. Instead, all cells uniformly displayed the same level of the active mark at the 3’ end of the cluster (**Fig. 4f**). This suggested that while a large proportion of adult OLGs present a primed repressive state at the HOX loci, a sub-set of OLGs with no H3K27me3 deposition at the 3’ end but high H3K27ac, may be capable of expressing these genes and could explain the low leaky RNA expression that was observed in single cell transcriptomic studies.

### Developmental architecture of HOX genes in iPS-derived human OLGs

HOX gene transcription has been shown to be regulated by the local chromatin architecture during early human development^62,63^. Given that we observed epigenetic memory of the developmental state of the chromatin at the HOX genes in adult OLGs, we questioned whether the 3D chromatin conformation plays a role in the epigenetic state of HOX genes in OLGs. The activation of the HOX genes in development is associated with the dissolution of a topologically associated domain (**TAD**) containing the HOX genes and the formation of two distinct centromeric and telomeric TADs (c-Dom, t-Dom) connecting the HOX genes with enhancers in the flanking regions^60,63^. To check the TAD structure in OLGs, we performed high depth Micro-C, a high-resolution proximity ligation-based assay for capturing genome-wide chromatin contacts^64^, in iPS-derived human OPCs^65,66^, capturing approximately 4 billion paired-end reads (**Extended Data Fig. 8a-b**). We also performed Micro-C in human primary memory B-cells, to compare with a terminally specified cell from another developmental lineage. We were able to capture broad compartment-level information^67^ as well as TAD structures^68^ at a resolution of 5kb, which corresponded with the well characterized *SOX9-KCNJ2* locus (**Extended Data Fig. 8a-e**). The active A compartments^67^ identified in the OPCs also corresponded to regions of high accessibility in OPCs, further strengthening the validity of the data (**Extended Data Fig. 8f**). Finally, we could also identify cell type-specific loops and interactions in both B-cells and the human OPCs (**Extended Data Fig. 8g**).

We checked the chromatin architecture around the HOXA and HOXD clusters and found distinct differences between hOPCs and B-cells. While in B-cells, these HOX clusters were tightly interacting within a TAD, in OPCs they displayed a more open architecture, and instead were interacting broadly with regions upstream and downstream of the clusters in two large TADs, reminiscent of the developmental c-Dom and t-Dom in development^63^ (**Fig. 5a**). A boundary analysis revealed a strong border within both clusters (**Fig. 5b**). We then overlaid the 3D architecture data with the accessibility and histone PTM data from the human adult spinal cord-derived OLGs and observed the TAD boundary between the c-Dom and t-Dom coinciding with the strong border identified earlier separating the primed from the silenced chromatin (**Fig. 5c; Extended Data Fig. 9a**). This suggested that the 3’ genes and 5’ genes of each cluster might be associated with an active and silent TAD respectively. Within the TADs, we also observed sub-contacts between regions outside the cluster. Interestingly, within the HOXA active TAD, we observed a contact with a region in the *SKAP2* locus, which contains a well-known enhancer regulating the expression of 3’ HOXA genes in development^69^ (**Fig. 5c**).

**Fig.5.**
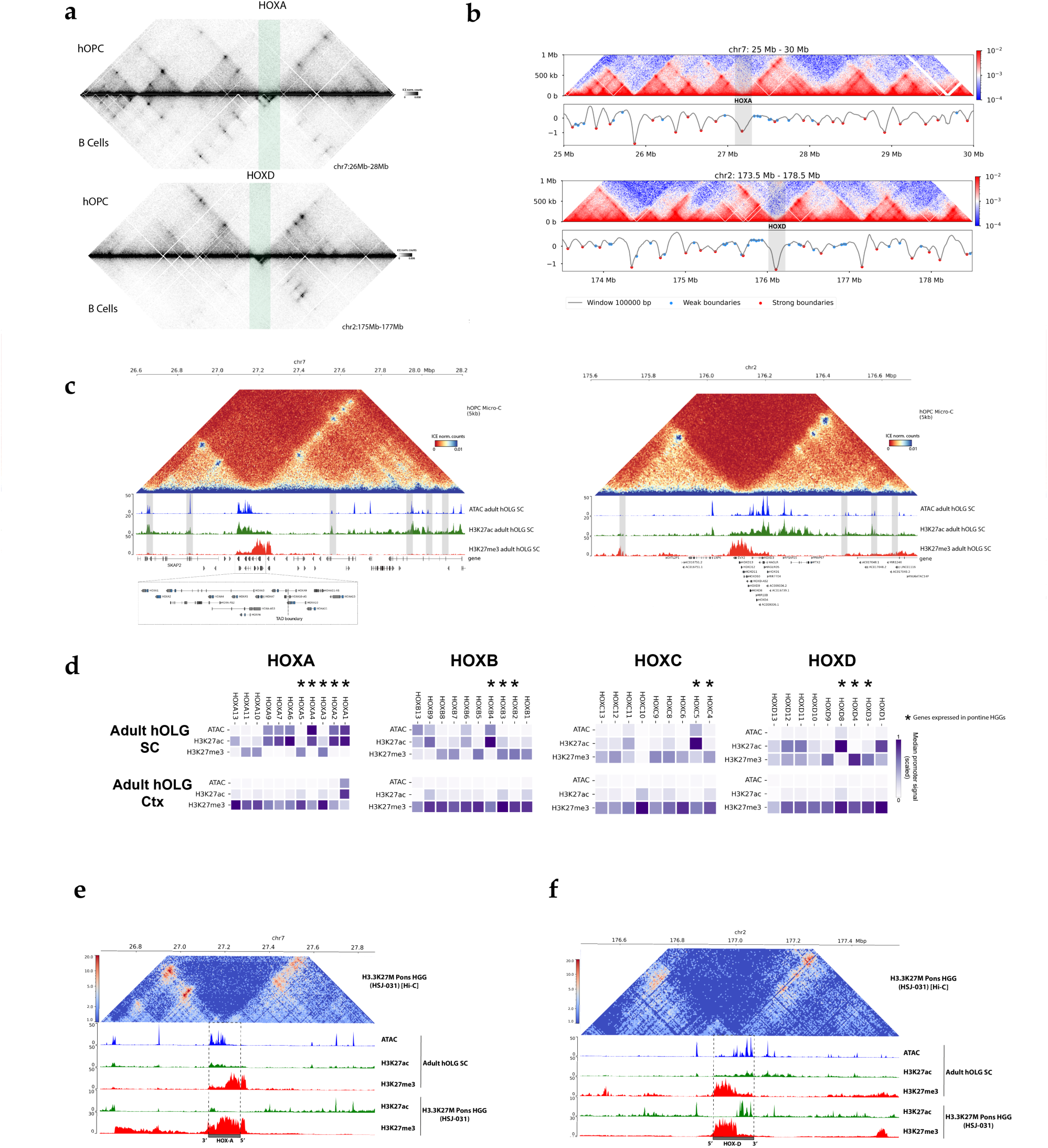
a. Normalized Micro-C contact matrix at 5kb resolution in iPS-derived hOPCs (upwards) and B-cells (inverted) at the HOXA (upper panel) and HOXD (lower) clusters showing the difference in HOX chromatin architecture in the two cell populations. b. Boundary analysis at HOXA (upper) and HOXD (lower) showing contact frequencies and the insulation scores under the matrix. Strong boundaries are marked in red. Location of HOXA and HOXD clusters are shown in grey. c. Normalized Micro-C contact matrix at 5kb resolution at HOXA (left) and HOXD (right) locus and corresponding ATAC, H3K27ac and H3K27me3 tracks in human adult spinal cord OLGs showing the c-Dom and t-Dom TAD structures including the sub-TAD contacts. Contacts with distal enhancers is shown by the grey bars. d. Normalized promoter accessibility (ATAC), H3K27ac and H3K27me3 signal in Spinal cord OLGs (top) and cortical OLGs (bottom) at all HOX genes. Asterisk marks the genes previously identified to be expressed in pontine high-grade gliomas (HGG). e. Normalized Hi-C contact matrix in H3.3K27M pontine HGG at the HOXA locus (marked by dotted lines) and corresponding ATAC, H3K27ac and H3K27me3 signal in spinal cord OLGs and H3K27ac and H3K27me3 in H3.3K27M pontine HGG, showing similarity in mark distribution in non-diseased conditions and gliomas. f. Same as in e) but at the HOXD locus.

### HOX genes with primed chromatin in spinal cord OLG are activated in high-grade gliomas

Ectopic activation of HOX genes is a feature of several cancers^60,70^. Within the CNS, midline high-grade gliomas (**HGG**) bearing the H3K27M mutation have been shown to have OPC origins^18,71^. These gliomas that affect children and young adults exhibit strong spatiotemporal specificity, arising from more posterior regions in the CNS, such as the thalamus, brain stem, cerebellum, and spinal cord^71,72^ (**Extended Data Fig. 9b**). They also present PRC2 function disruptions, affecting the global distribution of mono, di and tri methylation of the H3K27 residue^18^. The molecular architecture around the HOX genes in pontine and thalamic H3K27M HGGs faithfully recapitulates the locus of origin, providing a spatiotemporal address for the cell of origin^18^. Our observation of primed accessibility, histone PTM deposition and 3D chromatin architecture around several HOX genes in cervical spinal cord prompted us to ask whether the H3K27M-driven PRC2 disruption and specific HOX gene activation in midline/pons gliomas might be associated with the observed HOX priming in adult OLGs. Indeed, of the HOX genes that have been reported to be activated in pontine HGGs^18^, we found several promoters displaying the primed signature in spinal cord (**SC**) OLGs, but not in cortex OLG, where they display a repressed H3K27me3 associated state (**Fig. 5d)**. *HOXA1*, *HOXA3* and *HOXA5, HOXB4,* and *HOXD8* were all primed in SC OLG and expressed in posterior fossa group A ependymomas (PFA-EP) tumours and H3.1/H3.3K27M pontine HGG, but not more anterior H3.3K27M thalamic HGGs (**Fig. 5d; Extended Data Fig. 9c-f**).

We also overlaid the chromatin architecture and ChIP-seq data of H3.3K27M thalamus (more anterior) and pontine (more posterior, closer to the cervical spinal cord) HGG from a published dataset^18^. Strikingly, in pontine HGG we found a similar H3K27ac and H3K27me3 distribution pattern as well as the sub-TAD structures linking distinct HOX sub-clusters to remote enhancers, which were also primed with activation marks in human SC OLG (**Fig.5e-f**). In contrast, we did not observe overlap with thalamic gliomas, possibly due to the more rostral location of these tumours. Collectively, these findings suggest that the primed state of these genes in the non-diseased context in OLG in the posterior CNS may drive their expression upon PRC2 disruption in distinct brain tumours, being in line with the regional identity of the cell of origin in HGGs being a key determinant in their arisal^18^.

## Discussion

In this study, we profiled the single-cell chromatin landscape in adult human CNS from the primary motor cortex, cerebellum, and cervical spinal cord. We provide comprehensive datasets capturing the different cell types in the CNS at the level of chromatin accessibility (snATAC-seq) and histone-tail post-translational modifications (nanoCUT&Tag) in 108,626 and 58,696 cells, respectively. Our nanoCUT&Tag dataset in particular serves as a first-of-its-kind resource simultaneously capturing the H3K27ac and H3K27me3 landscape in single cells in different region of the adult human CNS. These unique trimodal epigenomic resources allowed us to identify and characterize a new candidate SOX10 enhancer specifically active in the OLG lineage. In addition, we provide a deeply sequenced chromatin architecture dataset in iPS-derived oligodendrocyte progenitor cells.

We identified significantly elevated chromatin accessibility in adult spinal cord, but not cortex, OLGs at the HOX cluster of genes (primarily in the HOXA and HOXD clusters), which was not correlated with robust gene expression. Profiling the histone landscape in these cells revealed a strong concordance between the ATAC and H3K27ac signal, and a negative correlation with H3K27me3. Nevertheless, while most of the active signal was located at the 3’ end of the clusters, there was a sharp inflection to a more repressed state on the 5’ end of the cluster. The elevated levels of H3K27me3 at the 3’ end relative to the regions immediately outside the cluster suggested that these genes may be kept in a primed state, as an epigenetic memory of developmental states. This memory was observed in the OLG lineage and astrocytes but not in microglia, possibly reflecting their distinct developmental origins. OLGs and astrocytes originate from the neuroectoderm, which undergoes the anterior-posterior patterning program involving HOX genes during development. In contrast, microglia derive from the yolk sac and migrate into the central nervous system after CNS patterning is complete, potentially explaining the absence of a similar pattern in microglia.

Epigenetic priming is commonly seen in pluripotent stem cells, where multiple fates are possible^73^. Genes associated with each lineage are kept both active and repressed simultaneously, to allow for rapid commitment and activation of a lineage and a quick repression of alternate lineages. Moreover, we have recently also shown that mouse OPCs can activate immune genes in the presence of an inflammatory insult and that these immune genes are also maintained in a primed state^74,75^. One reason for this might be to allow for rapid activation of immune genes to enable a quick response in an inflammatory context.

The role of HOX genes in development is well characterized and their expression in adult tissues occurs across several lineages^76,77^. However, the patterning identity of mouse oligodendrocyte lineage cells is strongly attenuated at postnatal stages, with *Hox* gene expression being actively downregulated^4^. Moreover, Hox gene expression in oligodendrocyte lineage cells in the juvenile mice is lower than in neurons^78^, suggesting different mechanisms of regulation of Hox genes in different neural lineages. In contrast, Hox genes activation is strongly correlated with various cancers^60^. Here, we show the preservation of chromatin architecture fidelity between our healthy spinal cord derived OLGs and H3.3K27M pontine paediatric high-grade glioma (HGG) tumour cells originating from OPCs. The priming of gene promoters, particularly those activated in pons HGG, coupled with the connections to distant enhancers, suggests that these genes are poised for activation in OLG from posterior regions of the adult CNS, but not in anterior regions as the cortex. The re-expression of these genes in homeostatic OLG is likely hindered by the deposition of H3K27me3. In turn thalamus HGG might require epigenetic memory at genes expressed developmentally at the anterior CNS, such as *OTX1*, *ZIC1*, *ZIC4*^18^.

Although this could explain a mechanism for HOX gene activation in pathological states, it does not reveal why epigenetic memory and priming is maintained to begin with. However, the HOX genes are also re-expressed in regenerative niches, and studies of limb injury have observed massive upregulation of HOX genes in regenerative stem cells^79^. A key feature of regenerative niches is the increased proliferation. OPCs are progenitor cells and more stem-cell like, and are also highly proliferative, especially in response to demyelinating injury^80^. Therefore, it may be possible that the HOX gene epigenetic priming may be associated with this proliferative potential. Indeed, spinal cord OPCs have also been shown to be more proliferative than those in the brain^81^, explaining why we find this signature more prominently in the spinal cord tissue. In addition, our network analysis shows the predicted HOX TF activity is higher in OPCs than in MOLs, further suggesting this regenerative potential may decrease upon differentiation. Thus, it is possible that one of the functions of the priming of these genes may be to allow for rapid gene activation in specific cellular contexts and in the context of remyelination and regeneration.

HOX epigenetic priming might also be deleterious, given the role of abnormal HOX expression in tumorigenesis. Within the active TAD of the HOXD cluster, we also observed a contact between *HOXD3* and *LINC01116*, which also contains an enhancer, which when activated induces hyper-proliferation of astrocytes in culture^82^ (**Extended Figure 9e,f**), a glioma-like phenotype^82^. Along with the 3D contact seen in the cultured OPCs, the spinal cord OLGs also displayed increased ATAC and H3K27ac signal at the LINC01116 locus. Interestingly, this increase in active signal is also seen in the cortical OLGs, although with heavier H3K27me3-mediated repression, suggesting that there may be specific contexts under which this pathway gets activated even in the brain. In the repressive TAD, we also observed long distance interactions with a distant element, which presented elevated levels of H3K27me3 deposition and lower levels of H3K27ac. Thus, our data suggests that 3D architecture at the HOX loci might regulate their transcriptional state in OLGs.

In conclusion, this comprehensive characterization of the epigenetic landscape of human adult neural cells, sheds light on how developmental epigenetic states might remain latent in adult cells, which could allow them to initiate regeneration processes, but also might prime them to unwanted transitions to tumour states.

### Limitations of the study

Our multiOme data gives the first insights into the single cell histone PTM landscapes of human neural cells in the adult CNS, in particular in the motor cortex and cervical spinal cord, revealing region-specific regulation of developmental genes. Our findings highlight that mapping of histone PTMs in different regions is relevant, and it would be of interest to investigate differences between further posterior CNS regions, such as thoracic and lumbar SC, and other anterior brain regions, such as the thalamus, hippocampus, among others. As our findings reveal developmental epigenetic memory relevant for high-grade glioma, probing other CNS regions might reveal additional disease susceptibilities. Our nanoCUT&Tag dataset exhibits increased sparsity compared to our previously published mouse dataset, though we attribute this disparity to the inherent challenges of working with frozen archival tissue, as opposed to fresh mouse tissue.

Our chromatin architecture data suggests that the HOXA and HOXD clusters may also be divided into two separate domains, which are reflective of the active chromatin architecture seen in development when these genes are being expressed. Interestingly, the strong recapitulation of the spatiotemporal context of HOX gene expression in our chromatin data suggests there might be a role for epigenetic memory, wherein the cells retain some memory of where they came from. Our Micro-C data has high resolution but is still acquired at the bulk level. Moreover, the human iPS-derived human OPCs are patterned for forebrain identity. Inducing a more posterior identity to iPS derived human OPCs or probing the chromatin architecture of human OLGs in different regions of the adult CNS could further elucidate how broad the mechanisms here described would be.

## Methods

### Human tissue collection and processing strategy

Adult post-mortem fresh-frozen tissue were obtained from the MRC Sudden Death Brain Bank in Edinburgh with full ethical approval (16/ES/0084). Work in Sweden was performed under the ethical permit 2016/589-31, with amendment 2019-01503, granted by the Swedish Ethical Review Authority (EPN). Tissue was collected from 20 different donors (10 male, 10 female) within the ages of 34 to 74 years old (**Supplementary Table 1**). Each donor donated fresh frozen white matter from the following three tissue regions: primary motor cortex (Brodmann area 4, BA4), arbor vitae cerebelli (CB) and fasciculi cuneatus and gracilis from and cervical spinal cord (CSC). Tissue was processed semi-randomly, ensuring that each batch of experiments had a representation of both sexes and all three tissue regions. Completed libraries were again randomly multiplexed during sequencing to minimize batch effects.

### Tissue dissociation and nuclei isolation

50-100mg of frozen tissue was placed in a 1.5mL tube and chilled in a mortar using liquid nitrogen. A chilled pestle was used to crush the tissue, followed by resuspension in 500uL nuclei permeabilization buffer (NPB; 5% BSA, 0.2% IGEPAL, 1mM DTT, 1x EDTA-free Protease inhibitor in PBS). Resuspended tissue was kept on ice for 15 minutes, with gentle pipetting every 5 minutes. The homogenized suspension was filtered through a 30um filter followed by a 10um filter. Equal volume of 50% Iodixanol solution was added and mixed thoroughly. 500uL 29% Iodixanol was gently underlaid using a syringe and needle, forming two phases. Samples were centrifuged at 4°C, 13,500xg, 20 minutes. Supernatant was removed and the nuclear pellet was resuspended in wash buffer (2% BSA in PBS). Samples were spun at 4°C, 1,000xg, 5 minutes. Supernatant was discarded and the pellet was resuspended in 30uL 1x diluted nuclei buffer (snATAC-seq; 10x Genomics) or 30uL 1x Antibody buffer (nanoCUT&Tag; recipe shown in corresponding section) or 30uL 1x diluted nuclei buffer with 1U/uL RNase inhibitor (multiOme; 10x Genomics)

### nanoTn5 purification and loading

Nanobody-Tn5 fusion proteins were purified as described earlier^83^. Purified enzyme was loaded using barcoded oligonucleotides. First, an equimolar mixture of 100uM Tn5_P5_MeA_BcdX_0N, Tn5_P5_MeA_BcdX_1N, Tn5_P5_MeA_BcdX_2N, and Tn5_P5_MeA_BcdX_3N were mixed with an equimolar amount of 100uM Tn5_Rev oligonucleotide. The oligonucleotide mixture was denatured by incubating at 95°C for 5 minutes in a thermocycler and allowed to anneal slowly by ramping down the temperature by 0.1°C/s. 8uL annealed oligonucleotide, 42uL glycerol, 44.1uL 2x dialysis buffer (100 mM HEPES-KOH pH 7.2, 200 mM NaCl, 0.2 mM EDTA, 2 mM DTT (freshly added), 0.2% Triton-X, 20% glycerol), 5.9uL anti-mouse nano-Tn5 (5 mg/mL, 67.7uM) or 8 µl annealed oligonucleotides, 42 µl glycerol, 45.7 µl 2× dialysis buffer, 4.3 uL anti-rabbit nano-Tn5 (6.8 mg/ml, 93 uM) to get a final 2uM of loaded nanoTn5 dimer.

### snATAC-seq library preparation and sequencing

Dissociated nuclei were counted and incubated at 37°C for 60 minutes in tagmentation mix. Tagmented nuclei were loaded onto the Chromium chip H (10x Genomics) according to manufacturer’s instructions. The Chromium Single Cell ATAC Library and Gel Bead Kit v1.1 (10x Genomics) was used to generate single-nuclei libraries. All libraries were sequenced on the Illumina NovaSeq 6000 with either the S Prime, S1, or S2 flow cell and a 50-8-16-49 read setup.

### nanoCUT&Tag

Multi-nano CUT&Tag libraries were prepared as described earlier. Briefly, dissociated nuclei in antibody buffer (20mM HEPES (pH7.5), 150mM NaCl, 0.05mM Spermidine, 1x Protease Inhibitor, 0.05% Digitonin, 0.01% IGEPAL, 2% BSA, 2mM EDTA in dH2O) were counted and 80,000-120,000 nuclei transferred to 0.5mL microfuge Eppendorf tubes. Nuclei were topped up to 96uL with antibody buffer. Mouse H3K27me3 antibody (1:100, Abcam #ab6002), rabbit H3K27ac antibody (1:100, Abcam #ab177178), barcoded anti-rabbit nano-Tn5 (1:100) and barcoded anti-mouse nano-Tn5 (1:100) were added to the nuclear suspension (final volume 100uL). Samples were then incubated overnight at 4°C on a rotator. After overnight incubation, cells were centrifuged at 600xg, 3 mins and washed twice with Dig-300 buffer (20mM HEPES (pH7.5), 300mM NaCl, 0.5mM Spermidine, 1x Protease Inhibitor, 0.05% Digitonin, 0.01% IGEPAL, 2% BSA in dH20). After the second wash, nuclei were resuspended in 100uL tagmentation buffer (20mM HEPES (pH7.5), 300mM NaCl, 0.5mM Spermidine, 1x Protease Inhibitor, 10mM MgCl2, 0.05% Digitonin, 0.01% IGEPAL, 2% BSA in dH20) and incubated at 37°C for 60 minutes. Tagmentation was stopped by adding 100uL STOP buffer (12.5mM EDTA in 1x diluted nuclei buffer (DNB, 10x Genomics) supplemented with 2% BSA). Nuclei were centrifuged at 600xg for 3 minutes and washed twice with 1x DNB/BSA to remove traces of EDTA. After the second wash, 185uL of supernatant was removed and nuclei were re-suspended in the remaining 15uL. 2uL was used for counting (1:5 diluted in Trypan Blue).

### nanoCUT&Tag library preparation and sequencing

Single-cell indexing was performed according to Chromium Next GEM Single Cell ATAC Library & Gel Bead Kit v1.1 (10x Genomics) instructions. 8uL nuclei was added to 7uL ATAC Buffer B (10x Genomics) and loaded onto the Chromium Chip H. GEM incubation and post-GEM incubation clean-up was performed according to Chromium Next GEM Single Cell ATAC Reagent Kits v1.1 instructions (Step 2.0 – Step 3.2). Of the 40uL of eluted sample, 2uL was used to measure the concentration using the Qubit dsDNA HS Assay kit. The remaining sample was used for P7 tagmentation by mixing with tagmentation mix: 2xTD buffer (20mM Tris (pH 7.5), 20% dimethylformamide, 10mM MgCl2), 1uL/10ng-template MeB-loaded standard Tn5, and dH2O up to a final volume of 100uL and incubating at 37°C for 30 minutes in a thermocycler. After tagmentation, samples were purified using DNA Clean and Concentrator-5 (Zymo) according to manufacturer’s instructions and eluted in 40uL Zymo elution buffer. Purified DNA was used as input for the Sample Index PCR in the Chromium Next GEM Single Cell ATAC Reagent Kits v1.1 (Step 4.1) and samples were amplified for 11-15 cycles. Post Sample Index Double-Sided Size Selection was performed according to manufacturer’s instructions. Library quality was checked on the Agilent bioanalyzer and sequenced on the Illumina NovaSeq 6000 S Prime flow cell (100c kit) with a custom read1 (R1_seq: 5’-GCGATCGAGGACGGCAGATGTGTATAAGAGACAG-3’) primer, custom index2 (I2_seq: 5’-CTGTCTCTTATACACATCTGCCGTCCTCGATCGC-3’) primer and a 36-8-48-36 read setup.

### multiOme library prep

Tissue dissociation and nuclei extraction was performed as described above. 10,000 nuclei were counted and used for bulk tagmentation followed by loading on the Chromium Next GEM Single Cell Chip J. Single-cell indexing and library preparation was performed using the Chromium Next GEM Single Cell MultiOme ATAC + Gene Expression kit, according to the manufacturer’s instructions. Libraries were sequenced on the Illumina NovaSeq 6000 S Prime flow cell (100c kit), with a 50-8-24-49 read setup.

### iPS-derived hOPC cell cultures

iPS-derived hOPC were derived in Steve Goldman’s lab, with the protocol described in Wang. et.al. (2013)^65^. Work in Sweden was performed under the ethical permit 2020-00398, with amendment 2023-04598-02, granted by the Swedish Ethical Review Authority (EPN). Corning 6-well cell culture plates were pre-coated with Poly-L-Ornithine (PLO, Sigma #P4957-50ML) and incubated for 1h at 37°C. PLO was removed and wells were rinsed three times using sterile 1x DPBS -/- (Thermo Fisher, #14190144) followed by overnight incubation with 5ug/mL Laminin (Corning, #354232) in HBSS +/+ (Thermo Fisher, #24020117). After removing the Laminin, 1 million iPS-derived hOPC cells (C27 line) were directly seeded into the plate and expanded for 3 weeks prior to splitting. Cells were cultured in proliferation media (DMEM/F12 (Invitrogen #11330-057) containing 1x B27 (Invitrogen #12587-010), 1x N1 (Sigma, #N6530), 1x NEAA (Invitrogen #11140-050), 60ng/mL T3 (Sigma, #T5516-1MG), 1uM dcAMP (Sigma, #D0260), 100ng/mL Biotin (Sigma, #B4639), 10ng/mL PDGF-AA (R&D #221-AA-50), 10ng/mL IGF-1 (R&D #291-G1-050), 10ng/mL NT3 (R&D #267-N3-025)), which was refreshed every 2 days.

### ATAC-seq in hOPCs

ATAC-seq was performed as described previously, with minor adaptations. 60,000 cultured hOPCs were collected, washed with 1x PBS, and incubated in lysis buffer (0.1% IGEPAL, 10mM Tris-HCl pH7.4, 10mM NaCl, 3mM MgCl2) on ice for 5 minutes. Lysed cells were centrifuged for 500xg at 4°C for 20 minutes. The nuclei pellet was resuspended in tagmentation mix (2xTD buffer, Tn5 enzyme, in dH2O) and incubated at 37°C for 30 minutes. Tagmented DNA was purified using the Qiagen minElute Purification kit and PAGE purified to remove adapter dimers. Libraries were sequenced on an Illumina NovaSeq 6000 with a 50-8-8-50 read setup.

### B-cell collection

Peripheral mononuclear cells (PBMCs) were freshly isolated by Ficoll (17-1440-03, GE healthcare) gradient centrifugation from buffy coats obtained through Karolinska University Hospital of 3 healthy donors female (aged 28, 29 and 39 years old). Study procedures were conducted under ethical permit 2009/2107-31-2 approved by the Swedish ethical review authority. B cells were then enriched by negative selection using EasySep™ Human B Cell Enrichment Kit II Without CD43 Depletion (17963, Stemcell Technologies) according to manufacturer’s instructions. B cells were then stained for 30min on ice with anti-CD3 (Clone SK7, 560176), -CD14 (Clone MφP9, 560180) and -CD16 APC-Cy7 (Clone 3G8, 560195), anti-CD19 APC (Clone HIB19, 561742), anti-IgG BV510 (Clone G18-145, 563247, BD Bioscience), anti-CD27 PE-Cy5.5 (Clone 0323, NBP1-43426, Novus Biologicals) or BV711 (Clone O323, 302833), anti-IgD Pacific Blue (Clone IA6-2, 348224), anti-IgM BV570 (Clone MMH-88, 314517) and Zombie NIR fixable viability dye (423106, Biolegend). Cells were then washed in PBS (D8537, Sigma-Aldrich) and filtered. Between 106 and 2×106 CD27+CD19+ Live B cells were sorted using a SH800 Cell Sorter (Sony) into a RPMI medium (R8758) with 10% heat-inactivated fetal bovine serum (F7524), 100 U/ml penicillin, and 100 µg/ml streptomycin (P4458, Sigma-Aldrich). Cells were washed in PBS, centrifuged, and stored at −80°C as dry pellet before proceeding with Micro-C.

### Micro-C

Micro-C was performed by the National Genomics Infrastructure, using the commercially available Micro-C kit (Dovetail Genomics, #21006). With 100,000-200,000 iPS-derived hOPCs as input. Briefly, cells were crosslinked and enzymatically digested using MNase to allow for nucleosome-resolution fragmentation. Free ends were ligated with biotin-containing adapters. Ligated fragments were reverse-crosslinked and amplified to introduce sequencing handles. Libraries were generated in three separate biological replicates and were sequenced as a pilot on the Illumina NovaSeq6000 S Prime flow cell with a 2×150bp (300c) kit with a 151-19-10-151 read setup. After quality control of the pilot and checking for library complexity (with the *preseq* tool), two of three hOPC replicates and all three B cell replicates were re-sequenced on a large S4 flow cell to a depth of 6 billion reads.

### snATAC-seq data pre-processing and QC

Fastq files generated from sequencing were processed on using “cellranger-atac count” with the default parameters. Samples were aggregated using the “cellranger-atac aggr” with default parameters, but with the normalization omitted using the flag “*—normalize=none*”. 2-kb count matrices were built using a custom script “build_large_mtx.py“which is a modified version of episcanpy’s^84,85^ build_atac_mtx.py script and allows for reading in files in batches. TSS enrichment scores were generated using the ArchR^86^ package and only cells with TSSe > 7 and number of unique fragments > 3000 were retained.

### snATAC-seq peak calling

The Fragments file (fragments.tsv.gz) was split according to celltype annotation and peaks were called using the *callpeak* function from MACS2^87^ with the following parameters: ‘*-f BED -g hs -q 0.05 –shift −100 –extsize 200 –nomodel –call-summits –keep-dup=1*’. Peak annotation was performed using the HOMER^88^ annotatePeaks.pl function and a custom GTF file with miRNA and snoRNA removed.

### snATAC-seq Downstream analysis

After cell filtering, the top 100,000 features were retained and normalization was performed using term frequency-inverse document frequency (TF-IDF), followed by singular vector decomposition (SVD). A nearest-neighbour graph was built in the lower dimensional space followed by leiden clustering. The highly variable features from each cluster were retained and used to repeat the TF-IDF, SVD, graph building and clustering for a total of 3 iterations. Batch correction was performed using Harmony^89^. Differentially accessible regions were identified using both the “rank_features” function in episcanpy v0.3.2 with Benjamini-Hochberg correction for multiple testing as well as the diffxpy (https://github.com/theislab/diffxpy/) package with Sex, Age and Tissue added as covariates.

### Gene activity matrix and cell type annotation

A gene activity matrix was built using a 5kb promoter region flanking the transcription start site of genes. The count matrix was smoothened using MAGIC^90^ with default parameters to improve the signal. The top 50 distinct differentially accessible genes for all cell types found in our published snRNA-seq dataset were used as cell-type metagenes and aggregate gene activity was calculated for all genes within each cell-type metagene, generating a metagene score for each cell. The metagene scores for the different cell types as well as individual marker genes were used to assign the clusters to the broad cell types. We could not identify spinal cord-derived neurons, though it is known these neurons are particularly sensitive and susceptible to hypoxia, possibly leading to difficulty in isolating them. An overwhelming proportion of all cerebellar cells in the dataset were composed of the CBEX cells (also known as cerebellar granular cells) These cells are tiny and densely packed within the granular layer of the cerebellum, close to the GM-WM border. We suspect the skewed distribution may have arisen from imprecise dissection when collecting the WM from the tissue.

### Integration with snRNA-seq data

The annotation of the cell types based on transcriptome data from external datasets was performed using two different references Jäkel & Agirre at al. 2019 and Seeker et al. 2023. Jäkel & Agirre et al. 2019 expression matrix was downloaded from Gene Expression Omnibus (GEO) repository with the accession number GSE118257, and then converted to Seurat^91^ v.4.3.0.1 object. Seeker et al. 2023 dataset was downloaded from https://cellxgene.cziscience.com/collections/9d63fcf1-5ca0-4006-8d8f-872f3327dbe9 as a Seurat object including all the cell types. scATAC h5ad file was converted to Seurat using SeuratDisk R library with Convert function with assay=”peaks” to h5Seurat format. Gene activities were calculated based on the chromatin accessibility signal using ENSEMBL genes annotations (EnsDb.Hsapiens.v86). For each annotated gene, the region of the promoter (500bp upstream the annotated transcription start site) and the gene body were considered (promoter+genebody). Gene activities were calculated using FeatureMatrix function from Signac^92^ v.1.10.0 with the promoter + genebody in GRanges format, from GenomicRanges^93^ library, and the cellranger-atac 2.0 fragments.tsv for all the samples. Then, they were added to the Seurat/Signac object as a new assay. Label transfer was performed using Seurat. The FindIntegrationAnchors function was applied to find anchors between scRNA reference dataset and the query scATAC gene activities. The FindTransferAnchors function was used to find transfer anchors from each of the references to the query, and then used to perform the label transfer with the TransferData function using canonical correlation analysis (CCA) with 20 dimensions.

### Motif analysis

For identifying TF motif differences, we used ChromVar, which incorporates per-cell normalization to correct for transposase bias and depth bias, to output a deviation score for each TF motif. Here, we used the version of ChromVar adapted in ArchR allowing scalability to large datasets. We applied the standard ArchR pipeline and calculated the deviation scores using the CIS-BP database as TF binding reference and then exported the motif deviation matrix to CSV format.

### Trimodal genomic clustering

10kb genomic bins spanning the hg38 genome were used as input peaks to the deeptools^94^ compute-matrix function. ATAC, H3K27ac and H3K27me3 bigwig files from each population were used to generate a matrix of normalized signal in each genomic bin. The mean signal across each bin across each modality and each cell type was used to hierarchically cluster the bins (rows) and celltype+modality (columns). Pearson correlation was used to identify the correlation between each column (identifying similar and dissimilar celltype+modalities based on whole genome patterns).

### Co-Accessibility analysis

Co-accessible regions were identified using Cicero^95^. Cell type specific pseudo-bulk bam files as well as single-cell count matrices were provided as input to identify pairs of genomic bins with increased co-accessibility across different cells of a population. Co-accessibility score cutoffs of 0.25 and 0.5 were used to identify significant interactions and high confidence interactions respectively.

### TF Regulatory network

The Core Regulatory Circuit (crc) package^96^ was used to identify the core TF network. Briefly, H3K27ac bam and bigwig files and Super Enhancers (via ROSE algorithm) for each cell population were used to scan for TF motifs within Super Enhancer regions using FIMO^97^. A network was built by inferring the number of interacting TF motifs in the proximal super enhancer of a TF. The TF strength was assigned based on the difference between the number of outbound edges (Regulated TFs) and inbound edges (Targeting TFs). A higher score suggests a TF has a stronger regulatory influence over other TFs.

### TF Footprinting analysis

Footprinting analysis was performed using the Regulatory Genomics Toolbox (RGT)^98^. Briefly, peaks were called for the different cell-types and the HMM-based Identification of Transcription factor footprints (HINT) framework was applied to identify active TF binding-sites using default parameters.

### Integration of multiOme-ATAC with snATAC-seq data

The ATAC object was subsetted to contain only the same donors in which the multiOme was performed. The ATAC and multiOme-ATAC 2kb objects were concatenated and normalized using TF-IDF, followed by SVD dimensionality reduction. A kNN classifier was used to annotate the cells derived from the multiOme-ATAC according to the closest neighbours found in the annotated ATAC object. Transferred annotations in the multiOme-ATAC object were checked using the RNA expression of marker genes in the multiOme-RNA object.

### nanoCUT&Tag data pre-processing and cell calling

Fastq files were split into antibody-specific fastq files using a custom *debarcoding.py* script. Once split, each antibody’s fastq file were used as input into 10x Genomics cellranger-atac v2.0 pipeline with the following parameters ‘*count –id=$(sample) –sample=$(sample) – fastqs=$(sample_fastqs) –reference=cellranger-atac/refdata-cellranger-atac-GRCh38-2020-A-2.0.0.*’ The output pseudobulk alignment file from cellranger was used to call peaks with MACS2 algorithm with the following parameters *‘callpeaks -g hs –keep-dup=1 –llocal 100000 –min-length 1000 –max-gap 1000’*. We then plotted the fraction of reads in peaks versus the total number of unique reads to custom select barcodes that we consider cells. The ‘is_cell_barcode’ bit in the singlecell.csv file from cellranger was reset or flipped to represent the called cells.

### nanoCUT&Tag peak calling and bigwig track generation

Peaks were called using the MACS2 *callpeak* function with the same parameters described above.

Fragments were split by cell type and used to generate a bam file with the bedtools bedtobam function. Cell type-specific bam files were sorted and indexed prior to bigwig generation. Bigwig files were generated using the following command – “*bamCoverage --normalizeUsing RPKM --binSize 50 --centerReads --smoothLength 250*”.

### Integration of H3K27ac nanoCUT&Tag with snATAC-seq data

The H3K27ac dataset was first subsetted to include only the cell barcodes shared with the H3K27me3 dataset. A new 2kb-count matrix was constructed for the H3K27ac dataset (query), and only features shared in the filtered ATAC dataset (reference) were retained. Cerebellar cells of the ATAC dataset were removed and the query dataset was integrated to the reference using the Scanpy ingest tool which is based on asymmetric mapping of the query data onto the reference’s nearest neighbour graph.

### nanoCUT&Tag signal enrichment

The k-means algorithm (k=10) implemented in deepTools v.3.5.1 was used to cluster all genes (1kb-padded TSS) based on the H3K27ac and H3K27me3 signal for each cell type. We then queried the genes identified in the clusters that displayed high H3K27ac and low H3K27me3 and using gget enrichr^99^, predicted the enriched celltype based on the genes (database=”celltypes”).

### Regulatory domain border identification

Normalized ATAC, H3K27ac and H3K27me3 signal distribution in genomic windows spanning 50kb upstream to 50kb downstream of each HOX cluster was calculated using deeptools compute-matrix. Each window was binned into 10kb bins, and the Kolmogorov-Smirnov test was used to identify pairs of adjacent bins with significantly different signal distribution for each modality separately. The shared border between the identified adjacent bins was considered as a “signal border”. Identified signal borders were annotated as weak, intermediate, or strong depending on if it was identified in only 1, 2 or all 3 modalities. Gaussian smoothening was applied to the signal for visualization in the plots, however all calculations were performed on the raw signal in each window.

### Micro-C data pre-processing

Micro-C data was analysed using the Dovetail Genomics analysis pipeline (github.com/dovetail-genomics/Micro-C/tree/main). The fastq files from the pilot and deep sequencing batch were merged and were aligned to the human genome (GRCh38) using bwa mem with the flags “*-5SP -T0 -t24*”. Pairtools was used to parse the aligned reads with the following command “*pairtools parse –min-mapq 40 –walks-policy 5unique –max-inter-align-gap 30 –nproc-in 16 –nproc-out 16*”. The count matrix in the .hic format was generated using the juicer package: “*java -Xmx32g -Djava.awt.headless=true -jar juicertools.jar pre –threads 16*”. Quality check of reads was performed using the get_qc.py script in the dovetail pipeline.

### HiC to Cool matrix conversion

.hic files are multi-resolution files and can be converted using *hic2cool* to generate multi-resolution .mcool files. However, due to the lack of updates in the hic2cool package, a workaround script “convertHic2Cool.py” was used which is an adaptation of the code sourced from (github.com/deeptools/HiCExplorer/issues/821#issuecomment-1316842070) and allows for generation of single resolution *.cool* files. Due to the space inefficiencies of storing multiple .cool files at all resolutions, individual cool files were generated from the parent .hic file for different analyses.

### Micro-C balancing and transformation

Raw contact matrices were normalized and balanced using iterative correction and eigen-decomposition (ICE) as implemented in the *cooler*^100^ package. The *hicTransform* package was used to generate O/E counts with the “*–method obs_exp*” flag.

### Insulation and boundary strength analysis

The *cooltools*^101^ python API was used to process the contact matrices and identify the insulation scores within the normalized contact frequency data. Briefly, a diamond-shaped window is used to slide along the genome, with one of the corners on the main diagonal of the contact matrix, and contacts within the window at each position are summed up. Windows with low sums are marked as putative boundaries and as insulating regions immediately upstream and downstream. Boundaries were identified in the 10-kb contact matrix with a sliding window size of 100kb.

### Loop calling and Virtual 4C analysis

Virtual-4C identifies loci that exhibit increased contact frequency with a reference locus of interest (viewpoint analysis) and was performed using the *hicPlotViewpoint* function in the HiCExplorer^102^ package. Loops were called on the 5kb contact matrix using the *mustache*^103^ package and looking within a maximum distance of 100Mb.

## Author Contributions

M.K and G.C.-B conceptualized the project and designed the experiments. L.A.S. performed macro-dissection of archival tissue blocks. M.K. and M.M. optimized the snATAC-seq protocol for archival tissue and M.K. optimized the nanoCUT&Tag protocol. M.K. performed single-cell experiments and analysed the data, with help from F.B.P (chromVar), E.A. (dataset integrations) J.Z and V.A. (data QC). K.C., N.R., M.J. and M.B. collected cells for and coordinated the Micro-C experiments with the National Genomics Infrastructure. M.K. analysed the Micro-C data. M.K. and G.C.B wrote the manuscript. All co-authors read and approved the manuscript.

## Acknowledgments

We thank Tony Jimenez-Beristain for writing the human ethical permit, Ana Pombo and Gioele La Manno for discussion, and Nada Jabado and Claudia Kleinman for critically evaluating the manuscript. We thank the donors, their families and the MRC Edinburgh Brain Bank for the archival tissue used in this study. We acknowledge support from the National Genomics Infrastructure in Stockholm funded by Science for Life Laboratory, the Knut and Alice Wallenberg Foundation, and the Swedish Research Council. Part of the computation/data handling was enabled by resources provided by the National Academic Infrastructure for Supercomputing in Sweden (NAISS) and Swedish National Infrastructure for Computing (SNIC) at the Uppsala Multidisciplinary Center for Advanced Computational Science, partially funded by the Swedish Research Council through grant agreement no. 2022-06725 and no. 2018-05973. Part of the computing was also performed in the Linnarsson group Monod Linux cluster at MBB-KI, and we thank Peter Lönnerberg for maintenance and support. Work in G.C.-B.’s research group was supported by the Swedish Research Council (grant 2019-01360), the European Union (Horizon 2020 Research and Innovation Programme/ European Research Council Advanced Grant SingleMS, grant agreement number 101096064), the Swedish Brain Foundation (FO2023-0032), the Swedish Cancer Society (Cancerfonden grant 23 2945 Pj 01 H), Knut and Alice Wallenberg Foundation (grant 2019-0089), the Göran Gustafsson Foundation for Research in Natural Sciences and Medicine, the Swedish Society for Medical Research (SSMF, grant JUB2019), Olav Thon Foundation, Ming Wai Lau Centre for Reparative Medicine, and Strategic Research Area Stem Cells and Regenerative Medicine (Karolinska Institutet). This project has been made possible in part by grant number 2019-002427 and 2021-239069 from the Chan Zuckerberg Initiative DAF, an advised fund of the Silicon Valley Community Foundation.

## Data & Code Availability

Raw Human data is currently being deposited in the European Genome-Phenome Archive (EGA) under EGA accession number EGAD50000000410. Browsable tracks are available at UCSC Genome Browser (https://cns-nanocuttag-atac.cells.ucsc.edu) and CZI CellxGene. All code is available at https://github.com/mkabbe/snATACnanoCT_AdultHumanCNS.

## Extended Data Figures

**Ext.Dat.Fig.1.**
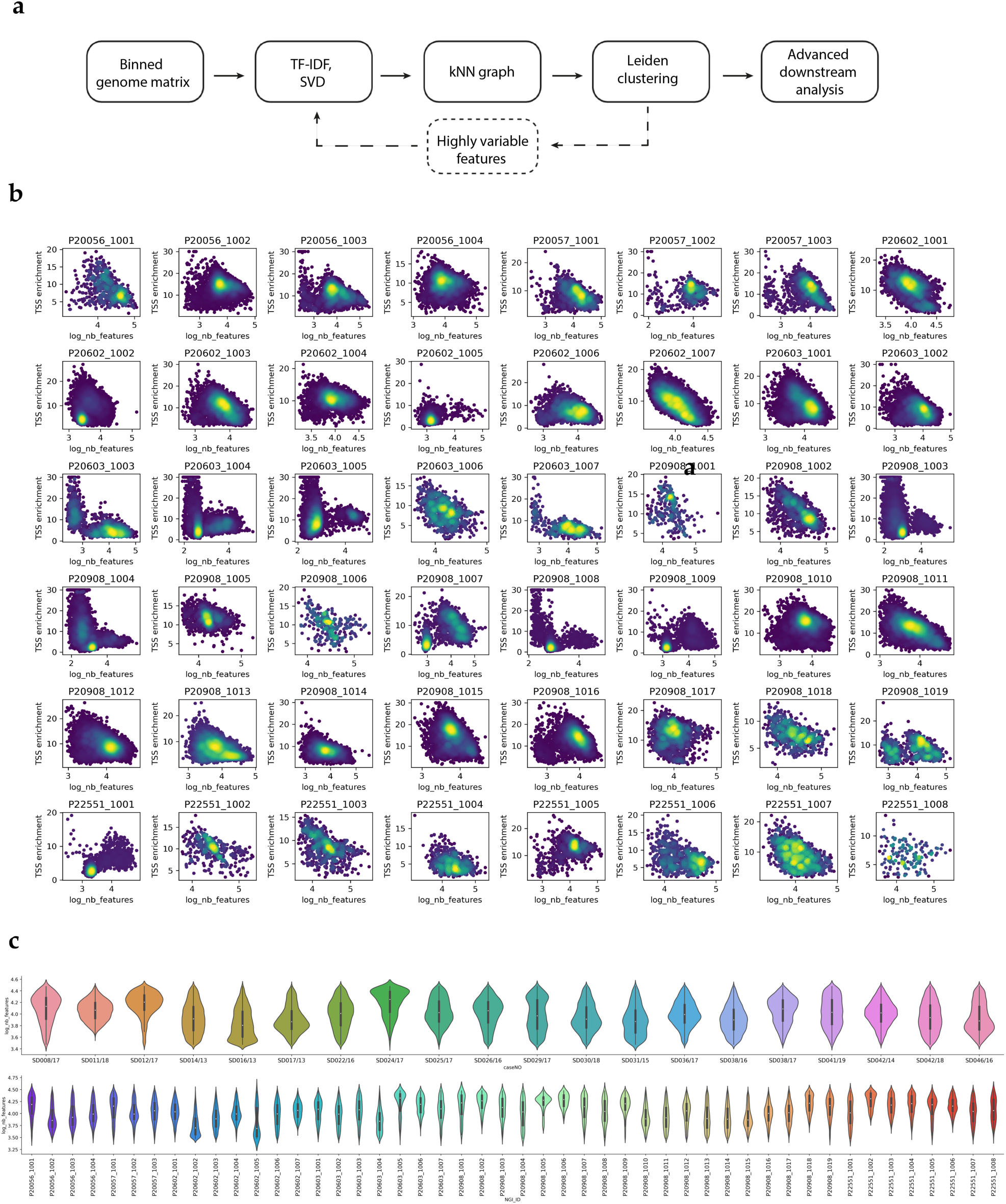
a. Schematic showing downstream data analysis workflow b. Density scatter QC plots for all 48 samples in the snATAC-seq dataset. Number of unique fragments (log scale) on x axis and TSS enrichment score on y axis. c. Violin plots showing number of unique fragments (log) from each donor. d. Same as c) but for each individual samples

**Ext.Dat.Fig.2.**
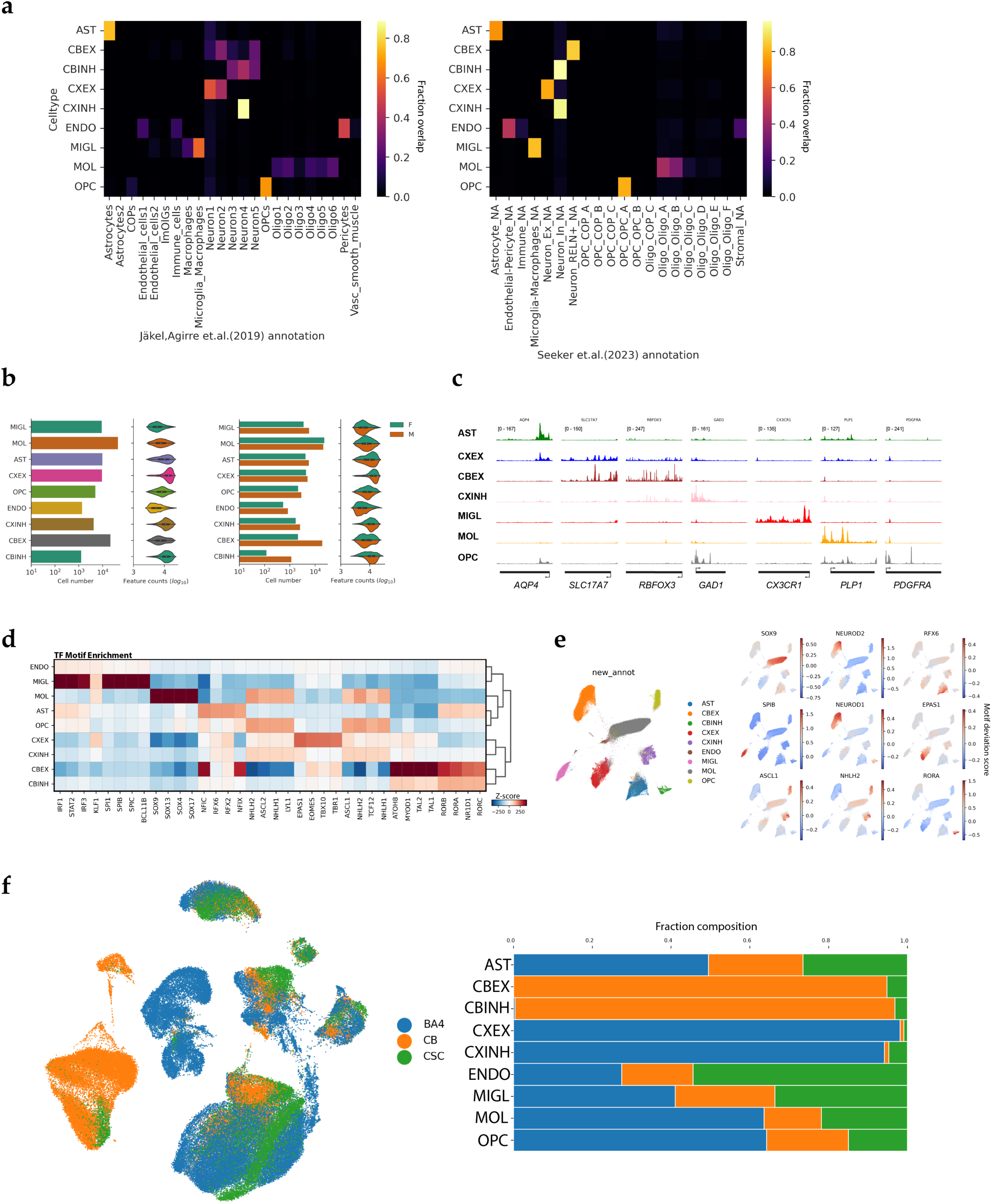
a. Correlation plot showing quality of celltype annotations using the metagene scores (y axis) and annotations from integration with previously published datasets (Jäkel, Agirre et.al 2019 – left, Seeker et.al. 2023 -right). b. Bar chart and violin plots showing cell numbers and feature counts (log scale) for each cell type (left) and contribution of each Sex to the metrics (right). c. Genome coverage tracks for different marker genes in each identified cell type. All columns are group normalized (range is shown in the first panel for each column). Gene location and TSS orientation is shown below. d. TF motif enrichment matrix showing the top 4 TF motifs identified as being differentially accessible in each cell type. e. 2D UMAP showing cell clustering based on the chromVar calculated TF motif deviations. Marker TF enrichment for each cell type is shown on the right. f. 2D UMAP (left) and bar chart (right) showing tissue composition of cell types in the snATAC-seq dataset.

**Ext.Dat.Fig.3.**
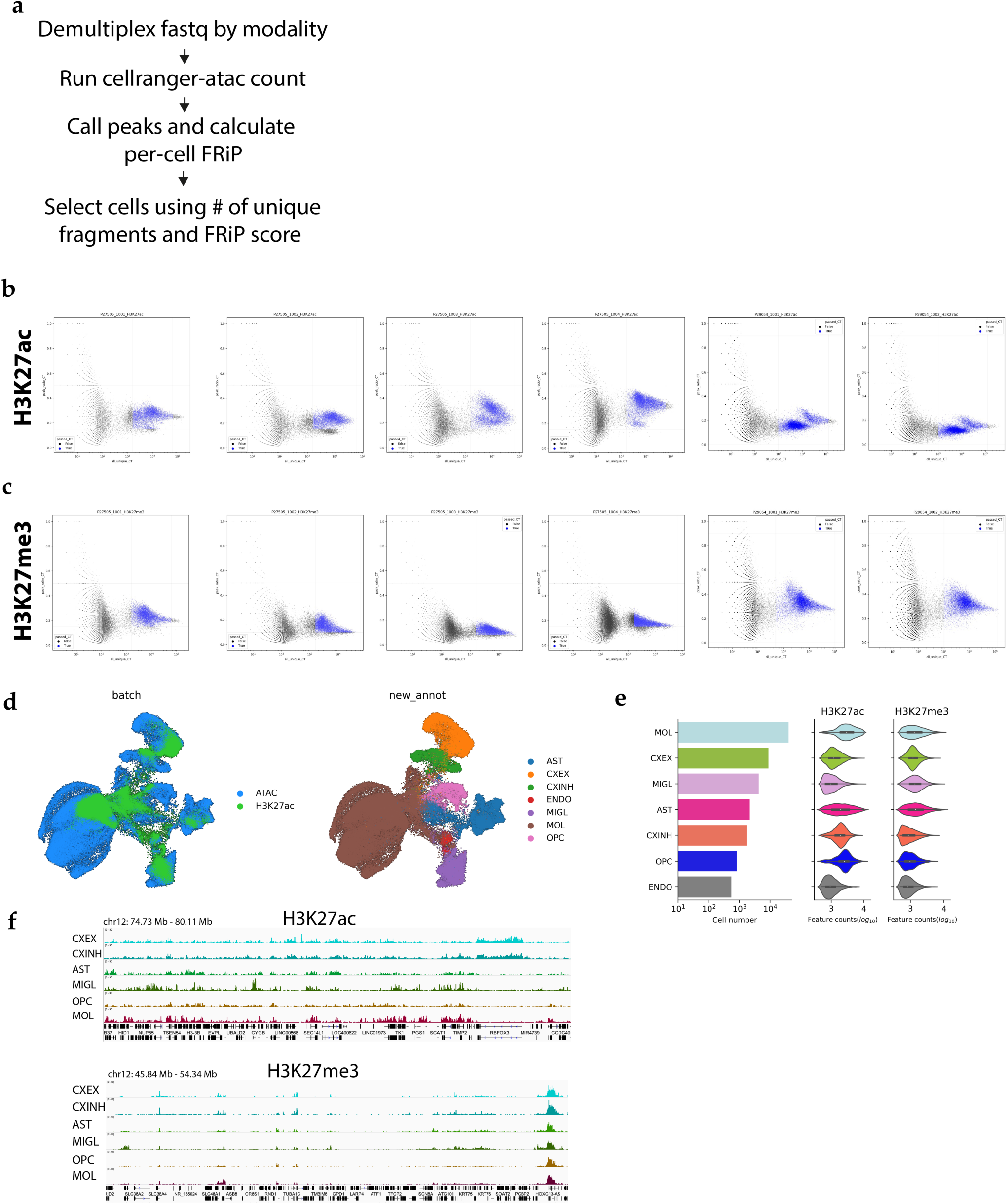
a. Workflow for demultiplexing and custom cell-calling for each nanoCUT&Tag antibody (H3K27ac, H3K27me3). b. Scatter plots used for custom cell calling in the H3K27ac modality. Number of unique fragments (log scale) shown on x axis and the fraction of reads in peaks (FRiP) shown on y axis. Selected cells shown in blue. c. Same as an b) but for H3K27me3 modality. d. 2D UMAP co-embedding of the ATAC and H3K27ac datasets, coloured by dataset (left) and cell population (right) e. Bar chart and violin plot showing the cell number in each population (left) and distribution of feature counts (log scale) for each modality (middle, right) f. Genome browser snapshot showing H3K27ac (top) and H3K27me3 (bottom) signal distribution across different loci for each cell type. Population specific signal in H3K27ac and pan coverage of HOX-C cluster with H3K27me3 can be seen.

**Ext.Dat.Fig.4.**
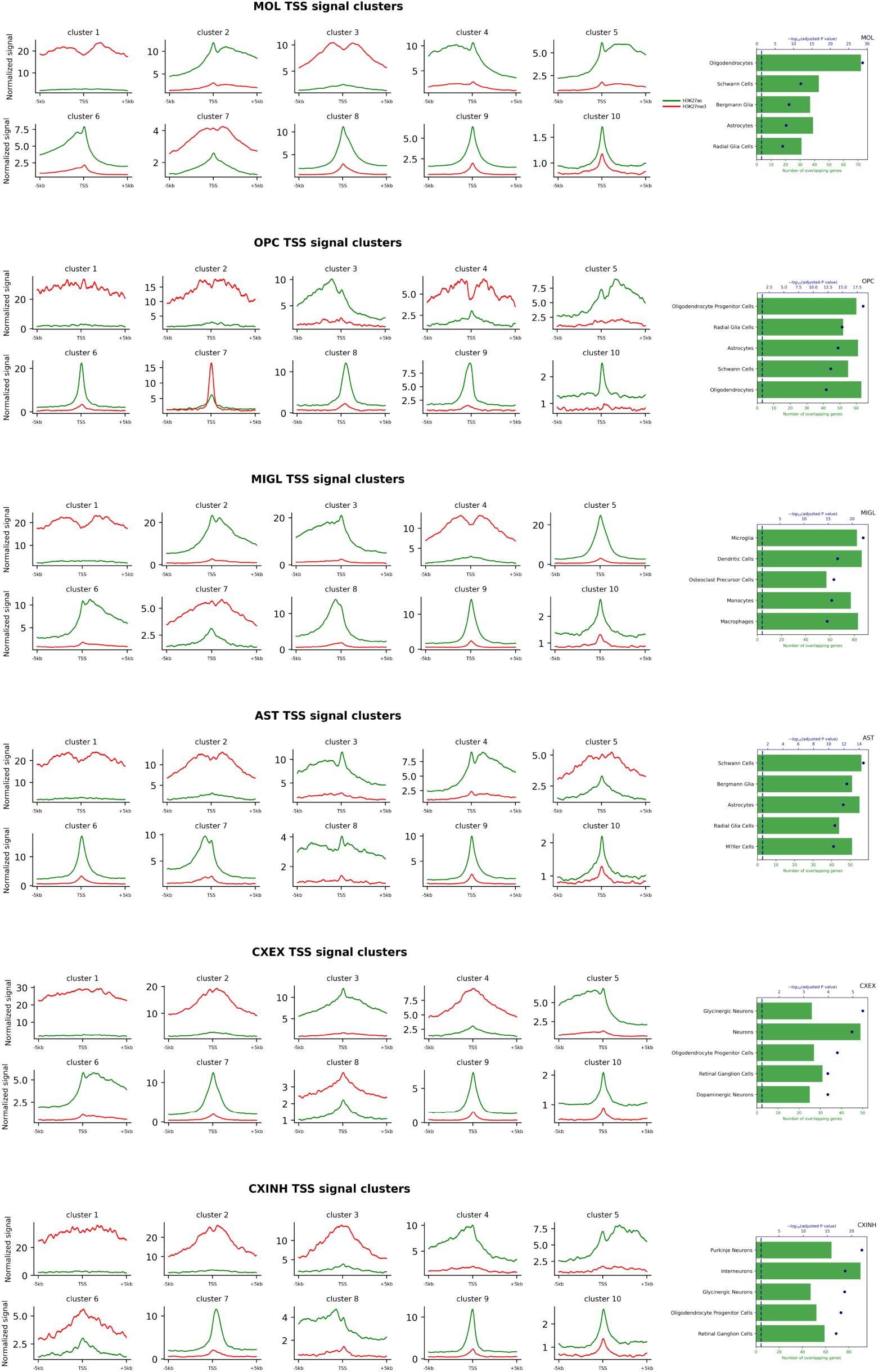
a. k-means clustering of H3K27me3 and H3K27ac signal distribution at the TSS of all genes for each cell type. Genes in the clusters showing strong H3K27ac and weak H3K27me3 were used as input for gget enrichr analysis to identify enriched cell types. Top identified cell type is the cell type itself highlighting the strong cell type specific signal captured in our nanoCUT&Tag dataset.

**Ext.Dat.Fig.5.**
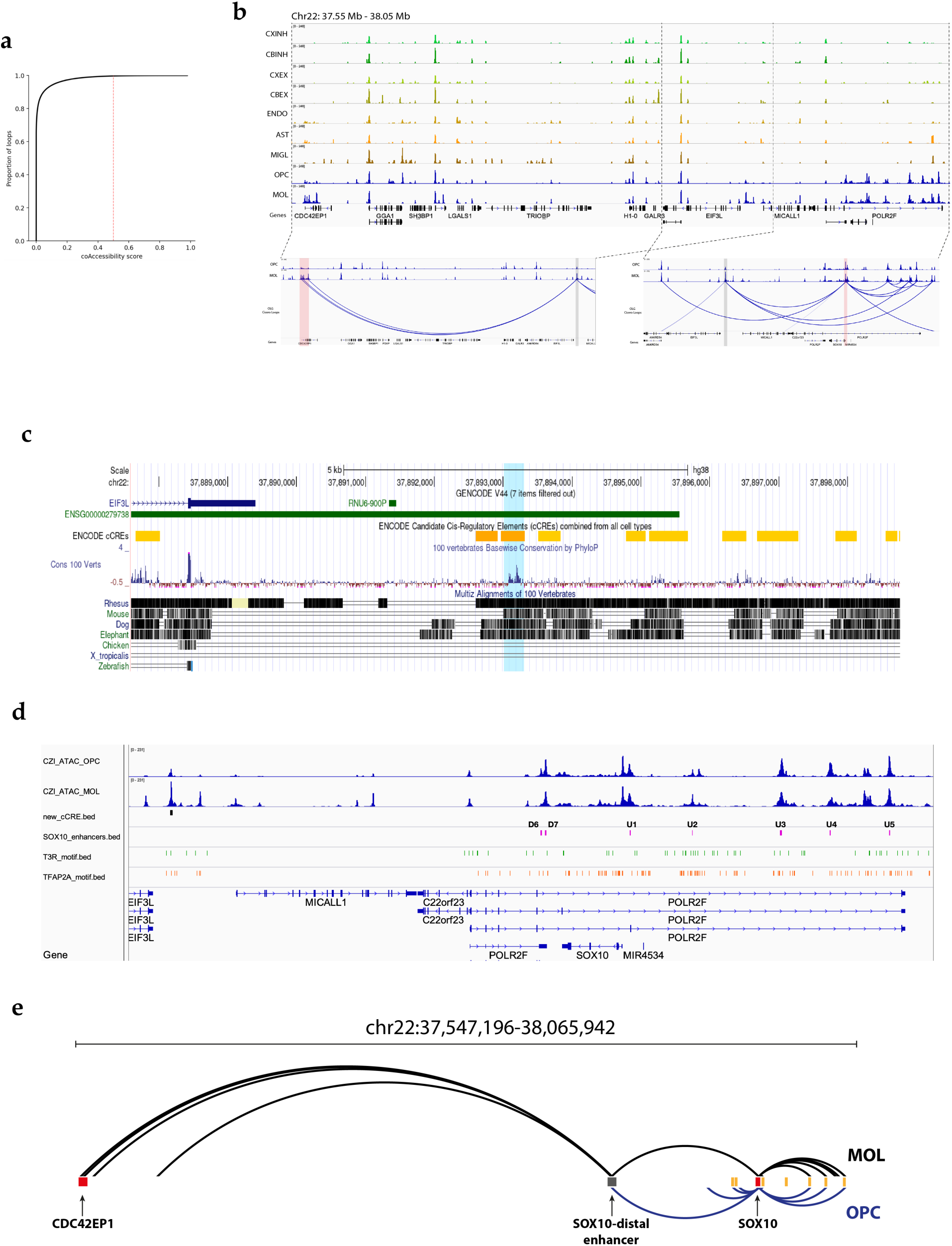
a. Cumulative distribution of the co-accessibility Score for all loops identified by cicero. Red line shows the score cut-off (0.5) used for assessing high quality interactions and captures the top 5% of all loops. b. Genome browser snapshot showing chromatin accessibility signal in all cell types at *CDC42EP1* locus (left), identified SOX10-distal enhancer (middle) and *SOX10* locus (right) and the corresponding loops identified using cicero. Red columns highlight the *CDC42EP1* and *SOX10* genes, grey column highlights the novel enhancer. c. UCSC Genome Browser snapshot showing the identified enhancer locus (light blue column) as well as overlap with previously identified ENCODE cCREs, PhyloP base conservation score and evolutionary conservation with different species. d. Genome browser track showing chromatin accessibility in OPCs and MOLs at the newly identified enhancer, SOX10 locus, and previously characterized U1-U5,D6 and D7 enhancers (purple) and overlap with thyroid hormone receptor motifs (T3R, green) and TFAP2A motifs (orange) e. Cicero loop comparison between MOLs (black) and OPCs (blue) showing shared loops with canonical enhancers (orange bars) and the newly identified enhancer (grey bar) and the MOL specific connection with CDC42EP1 (red bar)

**Ext.Dat.Fig.6.**
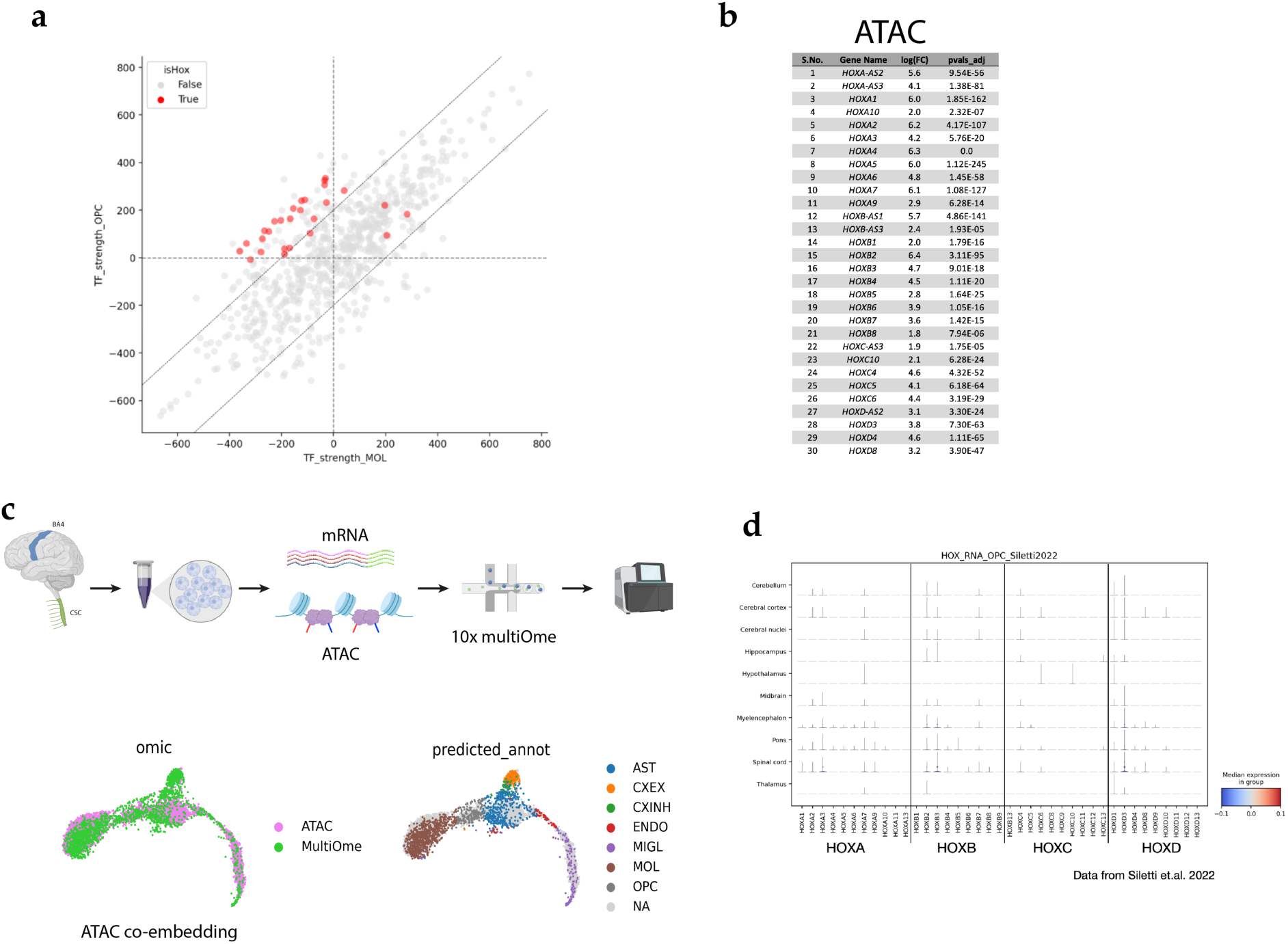
a. Scatter plot showing TF strength of shared core TFs in MOLs (x axis) and OPCs (y axis) and highlighting the identified HOX genes. b. List of identified HOX genes being differentially accessible in spinal cord OPCs and MOLs. c. Schematic showing workflow for the multiOme experiments, and cell type annotation using co-embedding followed by kNN clustering. d. Stacked violin plot showing gene expression levels of all HOX genes in OPCs in all regions from an adult human brain transcriptomic atlas (Siletti et.al 2023).

**Ext.Dat.Fig.7.**
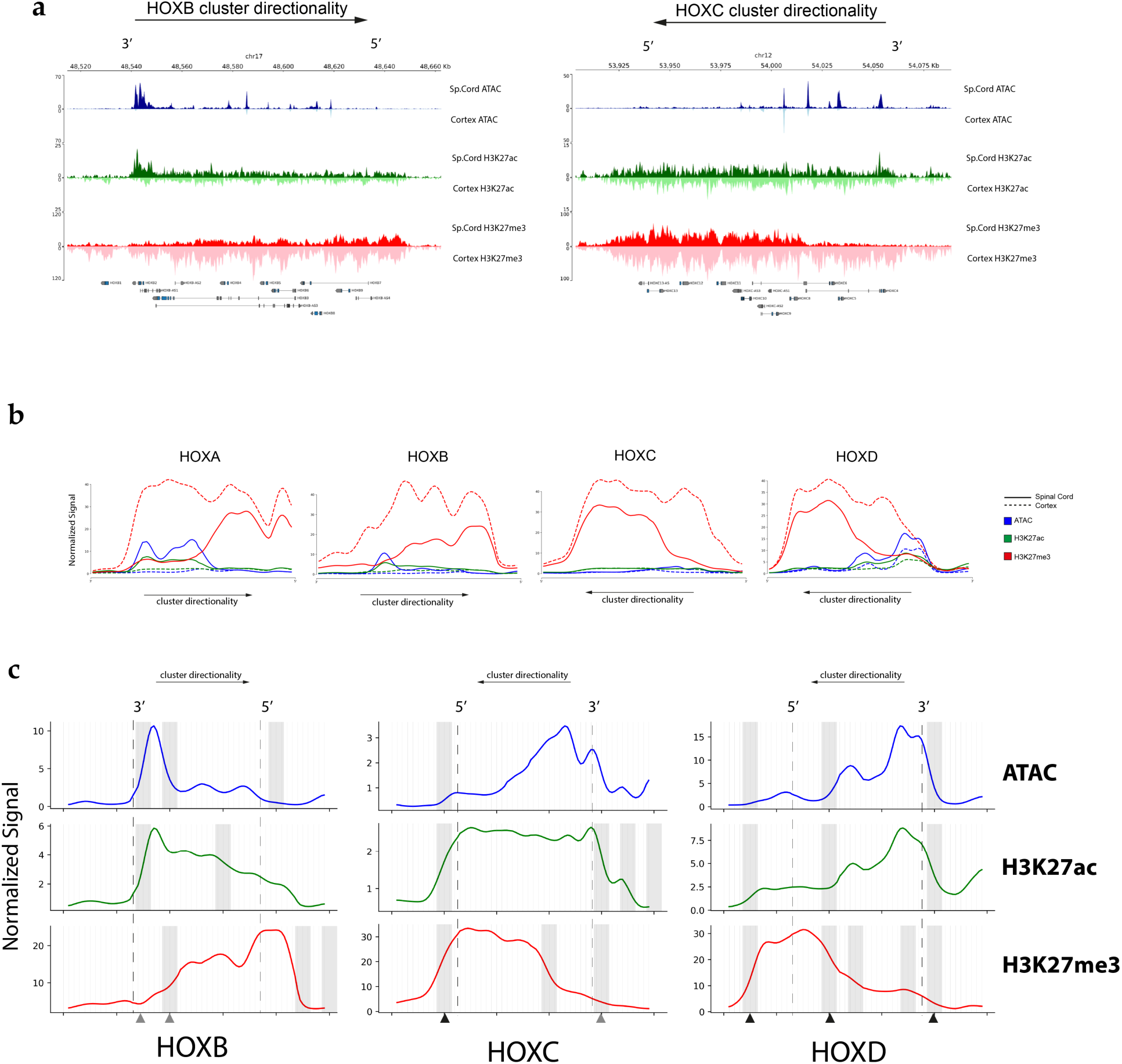
a. Genome browser tracks showing H3K27ac, H3K27me3 and ATAC signal in OLGs at the HOX-A, HOX-B, HOX-C and HOX-D clusters in cervical spinal cord (upright track, darker shade) and motor cortex (inverted track, lighter shade). Directionality of the clusters is shown by the arrow. b. Gaussian smoothed normalized signal from ATAC (blue), H3K27ac (green) and H3K27me3 (red) in Spinal cord OLGs (solid line) and cortical OLGs (dotted line) across each HOX cluster with a 50kb flanking region upstream and downstream. c. Gaussian smoothed normalized signal from ATAC (blue), H3K27ac (green) and H3K27me3 (red) across the HOXB, HOXC and HOXD clusters with a 50kb flanking region upstream and downstream, separated by each modality. Gray bars show the location of “signal boundaries” identified in each modality. Dotted lines mark the boundaries of each HOX cluster. Arrows underneath the plots mark intermediate and strong signal boundaries. HOX cluster directionality shown by arrow on top.

**Ext.Dat.Fig.8.**
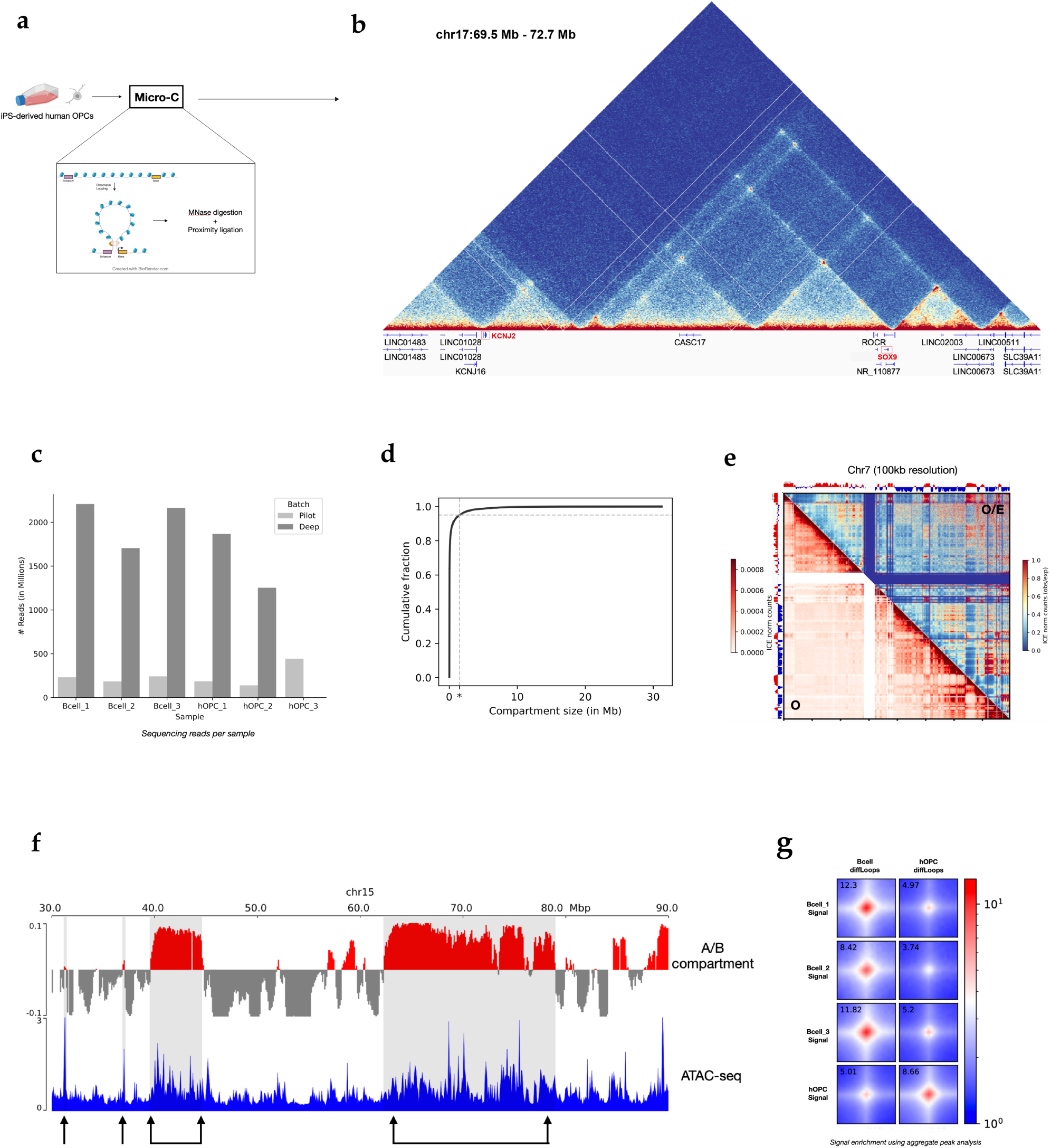
a. Experimental schematic showing collection of iPS-derived hOPCs and patient-derived B cells for Micro-C. Schematic of chromatin looping to bring enhancer and promoter in contact for transcription is shown below. b. Micro-C contact matrix in hOPCs at 5kb resolution showing the TAD formed at the *SOX9-KCNJ*2 locus. c. Bar chart showing the sequencing reads (in millions) obtained for each replicate in the B cells and hOPCs. Libraries were first shallow sequenced to assess library quality (light gray bars) followed by deep sequencing (dark grey). B cell replicates correspond to 3 separate patients. hOPC replicates correspond to separate biological replicates. d. Cumulative distribution of compartment size identified in the hOPC and B-cell Micro-C data. Asterisk marks 1.5Mb size as the upper size limit for 95% of all compartments. e. Contact matrix showing the normalized observed counts (lower triangle) and normalized obs/exp counts (upper triangle) at chromosome 7 in hOPCs. A and B compartments are shown along the sides of the matrix and exhibit strong corelation with “pockets” of increased contact frequency. f. A/B compartments in hOPCs on chromosome 15 overlaid on chromatin accessibility data from hOPCs (unpublished), showing corelation between active A compartments and increased accessibility. g. Pileup analysis of differentially accessible loops in B cells and hOPCs showing cell type specificity of identified loops.

**Ext.Dat.Fig.9.**
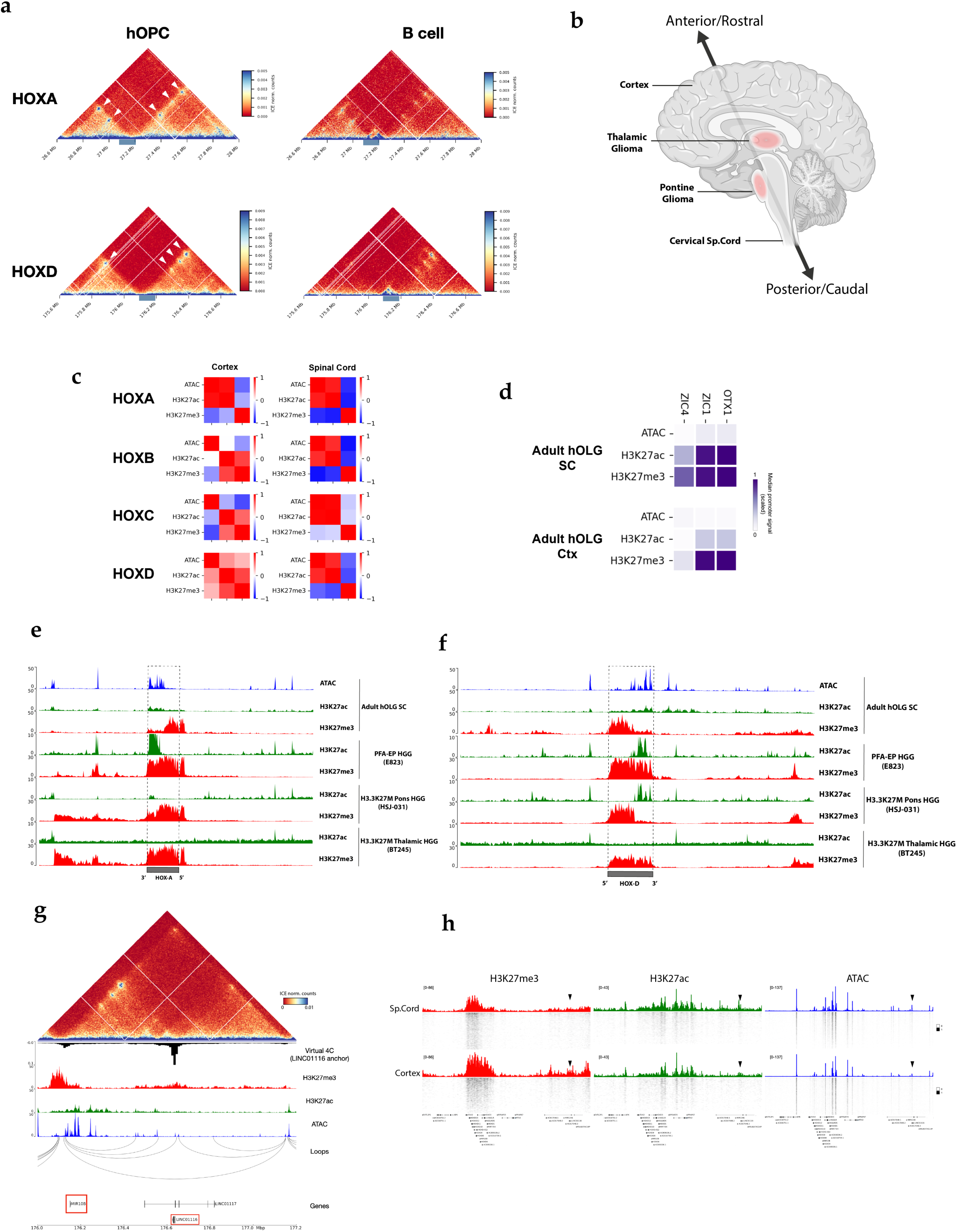
a. 5kb Contact matrix at HOXA (top) and HOXD (bottom) loci showing distinct TAD structures in hOPCs (left) and B cells (right). White arrows mark the increased contacts with distal enhancers in hOPCs. Location of HOXA and HOXD clusters shown by blue rectangle. b. Schematic of the adult human brain showing the location of pontine and thalamic gliomas along the A-P axis (created with BioRender.com). c. Correlation matrix of ATAC, H3K27ac and H3K27me3 signal across all 39 HOX promoters in Cortical (left) and Spinal Cord-derived OLGs (right), showing a stronger correlation between ATAC and H3K27ac in the spinal cord across all clusters. d. Normalized promoter accessibility (ATAC), H3K27ac and H3K27me3 signal in Spinal cord OLGs (top) and cortical OLGs (bottom) at the *OTX1*, *ZIC1*, and *ZIC4* genes which are relevant in formation of thalamic gliomas. e. Genome browser tracks showing H3K27ac and H3K27me3 signal coverage at the HOXA cluster in spinal cord (SC) derived adult human OLG (hOLG) and PFA-EP tumours, H3.3K27M pontine tumours, and H3.3K27M thalamic tumours. f. Same as c) but at the HOXD locus. g. Contact matrix showing long range interaction between MIR10-B and LINC01116, virtual 4c (anchored on LINC01116) H3K27me3, H3K27ac, ATAC and inferred loops are shown. h. Single cell ATAC, H3K27ac and H3K27me3 tracks showing signal distribution at HOXD and distal LINC01116 in spinal cord OLGs and cortex OLGs.

